# Deciphering Memory Patterns in Eco-Epidemiological systems through the lens of BaFOMS: A Bayesian Fractional Order Model Selection Method

**DOI:** 10.1101/2024.11.11.623021

**Authors:** Dipali Vasudev Mestry, Pratik Singh, Joacim Rocklöv, Amiya Ranjan Bhowmick

## Abstract

The evolution of eco-epidemiological systems is significantly influenced by the memory or previous history of the system. These non-Markovian dynamics are effectively modelled using fractional derivatives (FDs), incorporating memory kernels that reflect long-term or short-term memory characteristics in corresponding nonlinear fractional differential evolution equations. We introduce BaFOMS, a framework for identifying and selecting the optimal FD model for eco-epidemiological processes based on historical data. Specifically, we evaluate a class of fractional logistic growth models defined by their time correlation functions and determine the optimal model through Bayesian inference, selecting the one with the highest posterior probability. We also demonstrated BaFOMS efficiency in parameter estimation and forecasting, producing reliable results with quantified uncertainties. The method is shown to be robust across a range of eco-epidemiological datasets, offering computational efficiency and reliable inference about the evolution dynamics.

## Introduction

The concept of fractional derivatives (FD) dates back to 1695, when L’ Hospital and Leibniz discussed non-integer order derivatives as a generalization of integer-order calculus^1^. Since then, substantial advancements have elevated FD’s importance in mathematics, leading to innovative applications in engineering and natural sciences^1–7^. The inherent memory and hereditary characteristics, coupled with the increased degree of freedom, allows Fractional Differential Equations (FDE) to become an excellent tool in modelling real-world problems, particularly in epidemiology, ecology, and disease ecology^8–11^. These disciplines often involve nonlinear transmission dynamics and complex evolution and control processes, which are significantly influenced by past events and interventions. The incorporation of knowledge and historical context in these models are critical, as it profoundly shapes the evolution of models^12–15^. Classical integer-order ordinary differential equations (ODEs) models, characterized by Markovian or memoryless properties, may fall short in capturing the nuanced dynamics influenced by the historical trajectory of these systems due to their constant rate dynamics at all times^16^. While distributed time delay models attempts to introduce discrete memory by making the system’s current state depend on its state at a specific time in the past, this fixed time lag often leads to oscillations, instability, or chaotic behaviour especially for longer delays^17–20^. To address this limitation, fractional order models introduces a suitable continuous or discrete time correlation function that accounts for the effects of all past states of the system. This time correlation function can exhibit long-term memory decay, often following a power-law function, or short-term decay, similar to an exponential function. Such a framework offers a more comprehensive and accurate representation of the dynamic interplay between historical events and current system behaviour. This memory kernel is characterized by an explicit parameter that quantify the sensitivity (strong or weak) of the incorporated memory in system^9,20^. Additionally, FDE offer another leverage over ODE by providing the flexibility in incorporating memory from any point in the past, with recent states typically exerting more influence than distant ones, depending on the memory kernel. This flexibility means that if there is insufficient or inaccurate information regarding the initial conditions, the model can still evolve by introducing the initial effects at a more recent time^15^. Hence, the system becomes less dependent on precise knowledge of the initial conditions, enhancing the model’s resilience to uncertainties and inaccuracies.

To acknowledge quantitatively the fading effect of hysteresis memory, many definitions of FDs were proposed with suitable time correlation functions reflecting specific memory decay characteristics. The earliest formal FD definition, developed by Riemann and Liouville (RL) ^6^ 19th century, modelled memory as a power-law function. In 1863, Grunwald and Letnikov (GL) introduced an approach based on finite differences, later recognized as a numerical solution to the RL definition ^6,7^. A significant advancement occurred in 1967 when Caputo formulated an FD definition tailored for geophysical problems^21^, which is most widely applicable FD in eco-epidemiological modelling ^5,22^. Later, Caputo and Fabrizio (CF) ^23^ introduced a non-singular exponential kernel to address singularity issues, which inspired subsequent definitions, including Atangana–Baleanu-Caputo (ABC) ^24^, with a multi-parameter Mittag-Leffler kernel, and RL with exponential decay (CF-RL), proposed by Atangana and Aguilar^25^. Subsequently, many additional definitions were proposed, such as Hadamard ^26^, Riesz^7^ among others, each employing unique time correlation functions as memory kernels. Figure 1-A demonstrates the dynamics of a logistic growth model under most widely used FD formulations, including Caputo, CF, ABC, RL, CF-RL, and GL (Figure 1-B) all sharing identical parameters and initial conditions with memory parameter *α* restricted to *α* ∈ (0,1). A much more detailed and formal definition of these FDs can be found in Supplementary/S1.

**Figure 1.**
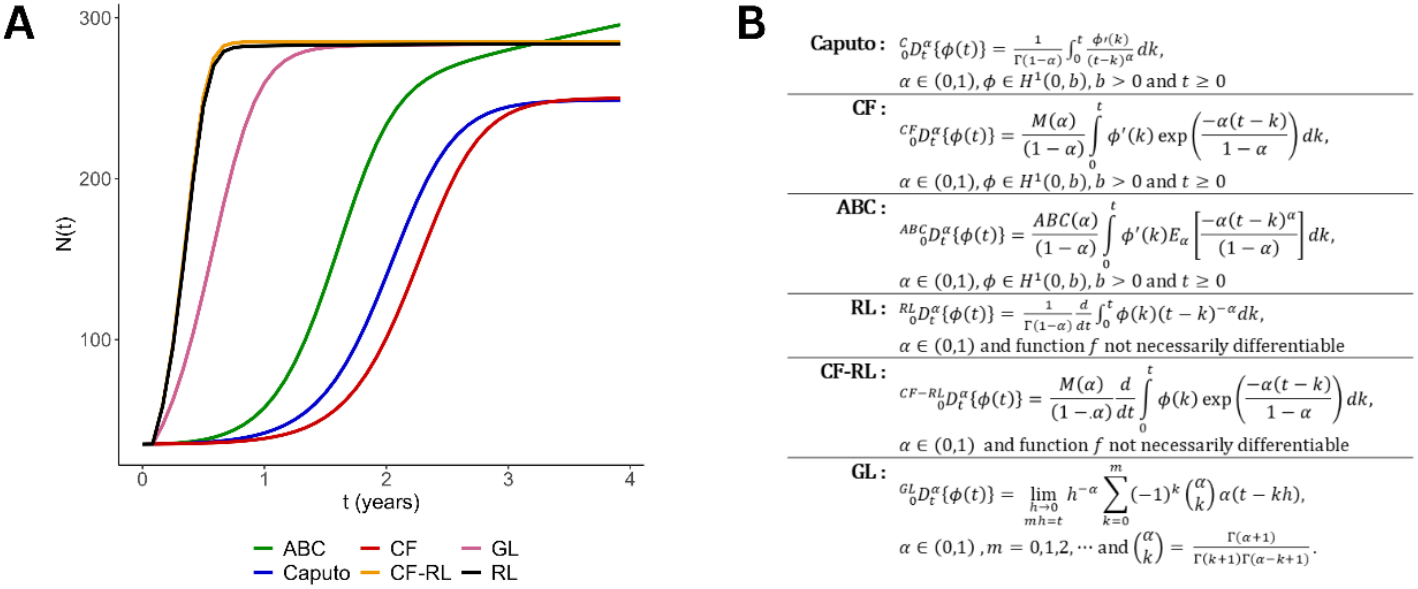
Fractional logistic growth equations Under Various FD. **A**. Solution curves for logistic growth equation under six FD definitions. The parameter values are kept as α = 0.001, r = 1.4, K = 1000 with an initial population size as N_0_ = 35. The step size of h = 0.001 and initial time point t_0_ = 0 were used. **B**. Definition of six widely used FDs in eco-epidemiological models. M(α), ABC(α) are the normalizing functions such that M(0) = M(1) = 1 and ABC(0) = ABC(1) = 1. The function E_α_(.) Is the generalised Mittag-Leffler function and Γ represents the standard gamma function.

Previous studies ^3–8^ have primarily examined the concept of FDs from a mathematical perspective, focusing on identifying equilibrium points, boundedness, existence, and uniqueness of solution space, bifurcation theory, and numerical simulation of the parameterized FD. Specifically, during Covid -19 pandemic there was huge surge of fractional dynamical models^10,27–30^ addressing both forward problems and inverse problems of parameter identifiability by fitting FDE models to data^16,24,25,31^. However, the choice of FD’s definitions in these studies, each associated with a specific memory kernel, often appears arbitrary, lacking any justification. The persisting question regarding the well-defined selection of a suitable FD definition prompts us to propose a method that leverages historical data of biological processes to quantify the associated memory within the system using Bayesian model selection framework. The growing availability of epidemiological, movement, migration, and population time series data offer an opportunity to analyse and quantify memory patterns in biological systems in a systematic way.

The proposed approach uses the Reversible Jump Markov Chain Monte Carlo (RJMCMC) ^32^ method to explore models with different parameter counts by allowing transitions between models of varying dimensions. This is further combined with a Gibbs sampling ^33^ technique to efficiently draw samples from the posterior distribution of model parameters and identify the model with the highest posterior probability, facilitating robust model selection. The goal of this method is not to definitively select the best FD among all available options, but rather to provide a framework for screening and determining which FD, with its specific memory kernel, is suitable for an eco-epidemiological system based on available historical data within defined levels of uncertainties. The selected FD will offer the most reliable inference on the dynamics of eco-epidemiological systems and along with optimizing computational efficiency. This systematic framework, in turn, has the potential to offer effective solutions to pertinent issues in public health, epidemic control, environmental management, and problems in the broad areas of mathematical biology. The improved accuracy and reliability of models derived from this approach enhance their utility in decision-making processes and contribute to advancing knowledge and solutions in these critical domains.

## Results

### BaFOMS architecture and plurality of fractional order models

To describe the BaFOMS architecture, we consider a simple logistic growth model with time dependent kernel as 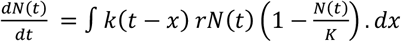, where *r* > 0, α ∈ (0,1], with initial condition as *N*(*t*_0_) = *N*_0_. The proper choice of time dependent kernel *k*(*t* − *x*) characterizes the plurality of the fractional order models. For instance, if we choose *k*(*t* − *x*) as power law correlation function as 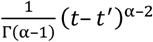, where 0 < α ≤ 1 and Γ(*x*) represents the gamma function, the right side of logistic growth model then is of the form R-L integral (Supplementary/Definition S1.3) and applying a fractional Caputo derivative of order *α* − 1 (Supplementary/Corollary S1.1) on both sides of model, we can easily recovered the well-known Caputo fractional order logistic equation. We propose a generic approach by taking six commonly studies FD’s (Fig 1-B) in eco-epidemiological processes and designate the logistic growth fractional model as *M*_1_ to *M*_6_. Each fractional differential model corresponds to a specific FD definition (Fig 1-B) and is restricted for *α* ∈ (0,1) i.e. *M*_1_ corresponds to Caputo, *M*_2_ to CF, *M*_3_ to ABC, *M*_4_ to RL, *M*_5_ to CF-RL, and *M*_6_ to GL. The models are then numerically discretized using suitable discretization schemes based on their memory kernel (Methods/Model development) to obtain approximate solutions. We further introduced gaussian noise in numerical solution (*μ*^(*m*)^(*t*_*i*_)) of each model to transform it into statistical models that accommodates uncertainty and given below:

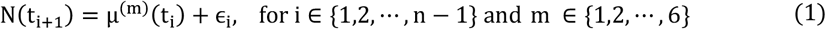

where ϵ_i_ ∼ 𝒩(0, σ^2^); and *N*(*t*_*i*+1_) denotes the (i + 1)^th^ observed value, incorporating uncertainty. The parameter *m* is introduced to distinguish among different fractional derivative (FD) definitions, ranging from *m* ∈ {1,2, ⋯, 6} representing Caputo, CF, ABC, RL, CF-RL and GL FDs definitions, respectively. The initial values of the state variable, N(t_0_) and *N*(*t*_1_), are subject to specific distributional assumptions as

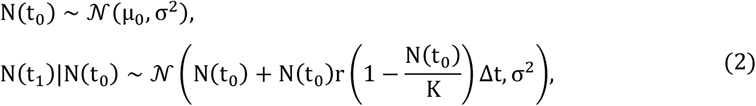

where *μ*_0_ is unknown and greater than 0. Hence, by utilizing equation (1) and (2)**Error! Reference source not found**., a unified expression for each FD definition can be formulated as follows:

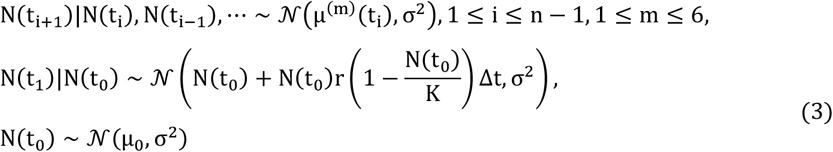

with the unknown parameter vector of interest θ = (α, r, K, σ^2^, μ_0_) ∈ θ = (0,1) × (0, ∞) × (0, ∞) × (0, ∞) × (0, ∞) ⊆ R^0^. It is essential to highlight that the distributions of *N*(*t*_1_), and *N*(*t*_i+1_), outlined in equation (2) and (3)**Error! Reference source not found**. are conditional over the previous population sizes. The order of the conditional dependence on the preceding time points is contingent on the approximate numerical solution *μ*^(*m*)^(*t*_*i*_) for *i* ∈ {1,2, ⋯, *n* − 1} and *m* ∈ {1,2, ⋯, 6}. The complete expression for *μ*^(*m*)^(*t*_*i*_), ∀ *i* and ∀m is given in (Supplementary/S2.4).

We then employed the Bayesian framework to compute the posterior distribution by evaluating data likelihood and assumed prior distribution of parameters. The conditional likelihood function is computed using Markov property as model’s present state is dependent on previous two preceding time points (***M***_**1**_ − ***M***_**3**_) or all preceding time points (***M***_**4**_ − ***M***_**6**_) (Methods/Likelihood function). The posterior probability of each model given the data can be calculated using the Bayes factor. Bayes factor^34,35^ for comparing model *M*_*i*_ with model *M*_*j*_ is given as

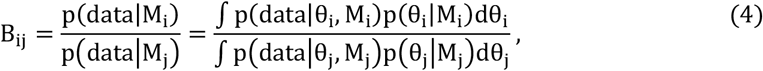

where p(data|θ_m_, M_m_) is the joint probability density for data and p(θ_m_|M_m_)is the prior distribution of model parameter *θ*_1_ for model corresponds to m ∈ {i, j}. Hence, computed posterior model probabilities using the Bayes factor is

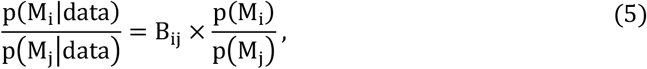

where p(M_m_|data) is the posterior model probability and p(M_m_) is prior model probability of m^th^ model. The computation of posterior model probabilities, involving complex integrals over a multidimensional parameter space, was conducted using a restricted version of the Reversible Jump Markov Chain Monte Carlo (RJMCMC) method elucidated by Barker and Link^36^. For implementation of RJMCMC in ***R*** software through ***rjmcmc*** package^35^, we utilized the Just Another Gibbs Sampler (JAGS)^33^ via the ***R2jags***^37^ package, to compute the posterior distributions of model parameters, enabling RJMCMC simulations to estimate posterior model probabilities. The most probable definition of FD’s is the equation having highest posterior model probability. The overall workflow for fractional order model selection (BaFOMS), is summarised in Fig 2-A and detailed architecture of numerical discretization and Bayesian framework is given in Methods and Supplementary/S2.

**Figure 2.**
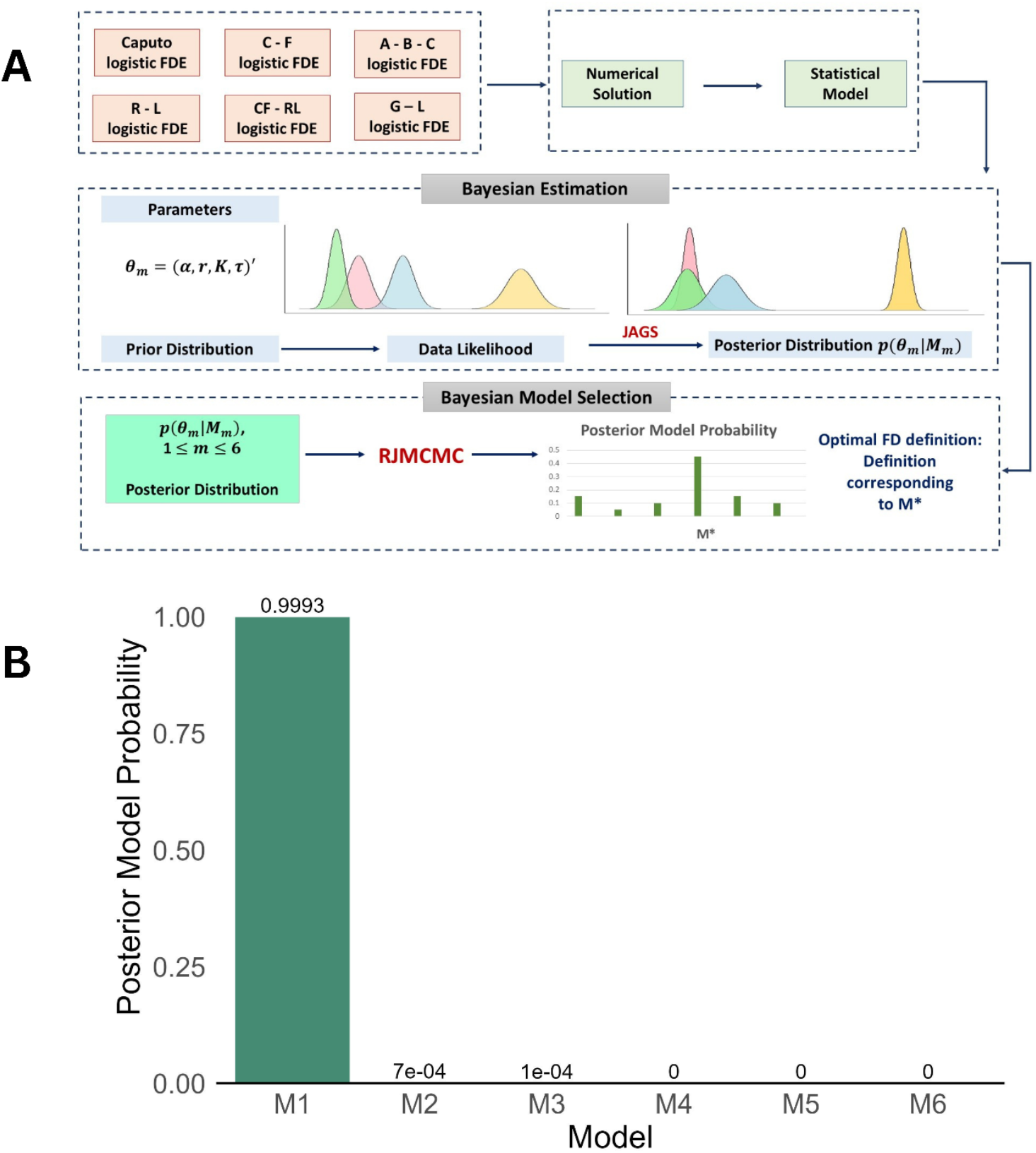
BaFOMS workflow and model evaluation Results. **A**. Schematic representation of the complete workflow, including different steps of analysis. Six distinct definitions of FDs are considered for the study and leading to the formulation of six corresponding logistic FDES. Numerical solutions for these equations were computed using various numerical methods. Subsequently, noise was introduced to these solutions to construct related statistical models. Within the Bayesian framework, posterior distributions of the model parameters were derived using JAGS and then model selection was performed using the RJMCMC method. The FD associated with the highest posterior model probability was identified as the optimal choice. **B**. Posterior model probabilities for each FD models within BaFOMS architecture for simulated dataset. The simulated dataset was generated using parameter settings (α, r, K, σ^2^, μ_0_)^′^ = (0.3,1.1,30000, 300^2^, 5000)^′^ over the time interval t ∈ [0, 20] from the model M_1_ (Caputo FD).

To further emphasizes on pluralization of fractional order derivatives and their different nature in capturing memory dynamics in eco - epidemiological processes, we captured the model predicted uncertainty in the observed data of three ecological (Population time series of two mammalian species *Ursus Americanus* (GPDD ID: 116), *Castor canadensis* (GPDD ID: 200) and one avian species *Phalacrocorax carbo* (GPDD ID: 9330))^38^ (Methods/Ecological population data) and Covid-19 epidemiological dataset of Germany^39^ ((Methods/Covid-19 data) following logistic growth curve using BaFOMS. We sampled parameter values from the joint posterior distribution of the parameters (obtained from JAGS) and simulated the population sizes in the given time window. The process has been repeated 1000 times and at each time point, the 95% confidence interval (CI) is obtained as (*N*_0.025_, *N*_0.975_), where *N*_0.025_ and *N*_0.975_ are the 2.5^th^ and 97.5^th^ percentiles of the simulated population sizes (Fig 3 and Fig 4). Fig 3 and Fig 4 clearly depict the pluralization of memory kernel and choice of proper FD before making inference on eco-epidemiological processes and our BaFOMS approach offers a promising way to achieve this goal. Models *M*_3_, *M*_4_, and *M*_5_ (Fig 4) failed to generate the total cases values using the sampled parameters values from their joint posterior distribution in case of Covid-19 dataset of Germany. The posterior probability of these models driven to zero as RJMCMC explores model space and identifies that these definitions provide little to no support given the data structure and parameter priors.

**Figure 3.**
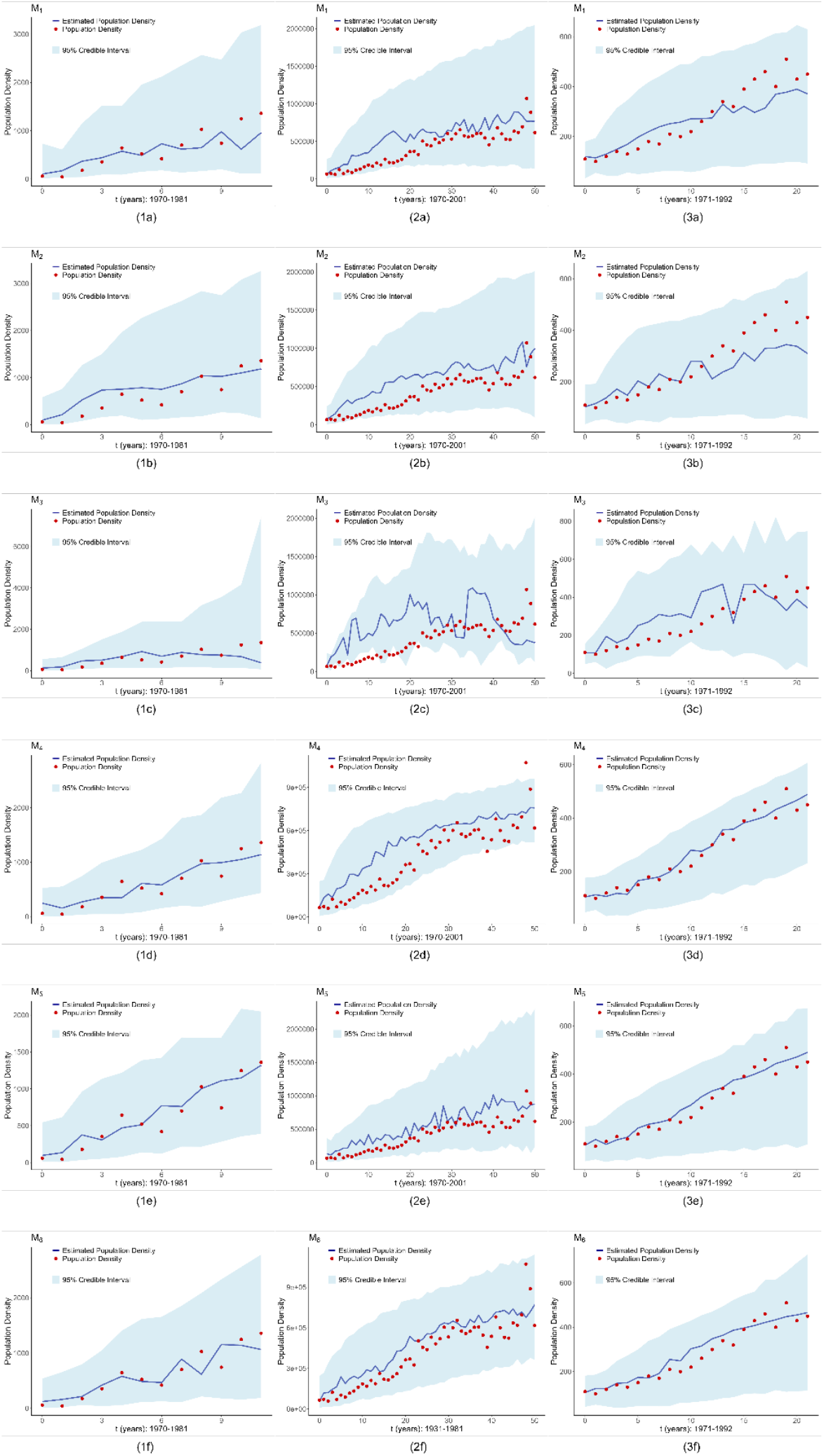
Fractional order model performance on ecological datasets. Each column presents six plots based on models on six models M_1_ to M_6_ for single-species analysis. The first, second, and third columns correspond to the species Ursus americanus (GPDD ID: 116), Castor canadensis (GPDD ID: 200), and Phalacrocorax carbo (GPDD ID: 9330), respectively. Parameter values were sampled from the joint posterior distribution obtained using JAGS, and population density values were generated over time. This process was repeated 1000 times to calculate the 95% confidence intervals (CIs). The mode of the simulated population size at each time point was taken as the predicted population size (blue colour line). The red solid points represent the data points, the blue line indicates the fitted values, and the light blue ribbon depicts the 95% CI

**Figure 4.**
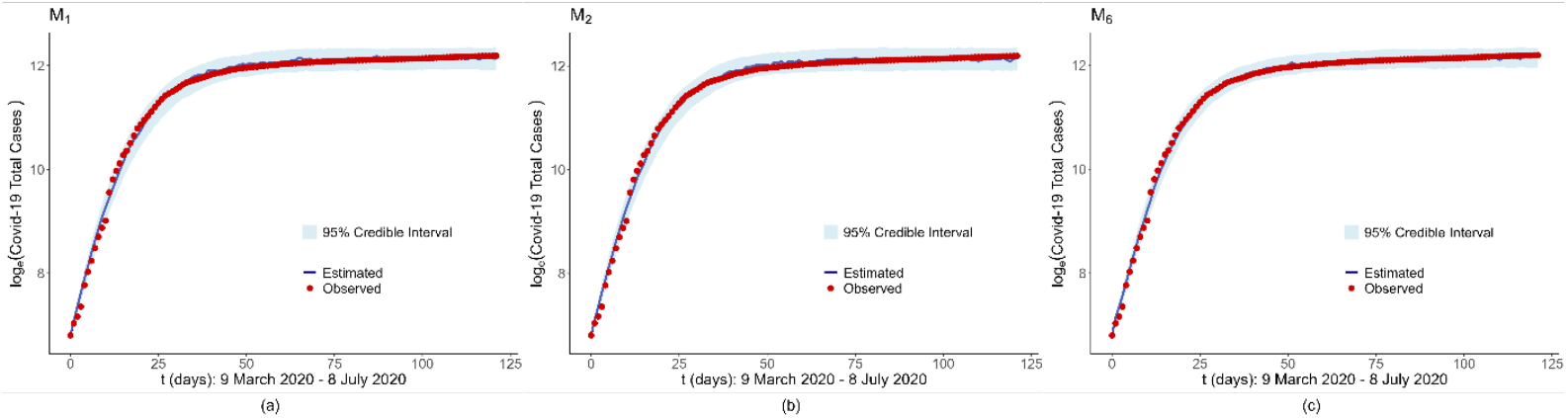
Fractional order model performance on epidemiological datasets. Epidemiological dataset (Covid-19) of Germany is analysed. Parameter values were sampled from the joint posterior distribution obtained through JAGS, and total case values were simulated over time. This process was repeated 1000 times, and the 95% confidence interval (CI) was calculated. The mode of the generated values at each time point was taken as the fitted total case value. Red solid points represent the observed data, the blue line shows the fitted values, and the light blue ribbon represents the 95% CI. The plots (a), (b), and (c) correspond to models M_1_, M_2_, and M_6_ (representing Caputo, CF, and GL FD definitions, respectively). Other models did not generate total case values successfully using parameters sampled from their joint posterior distributions, and hence, their plots are not shown.

### BaFOMS Performance Assessment and Model Selection

To validate the effectiveness and robustness of our approach, we simulated datasets using Caputo FD (model *M*_1_) of length 40 (n = 40) for t ∈ [0, 20]. The parameter values of model are kept fixed as (α, r, K, σ^2^, μ_0_) = (0.3, 1.1, 30000, 300^2^, 5000) for simulation study. Following this, we computed the posterior probabilities for each FD’s model mimicking the generated dataset using BaFOMS approach. In model fitting, prior distribution of parameters of each FD model are assumed to be independent and considered as follows:

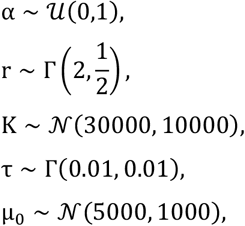

where precision parameter 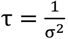. Since the model parameters are assumed to be a priori independent, prior distribution of π(θ) is

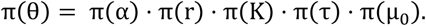

Since each FD model have same parameter dimension equal to 5, the common initial values for all the parameters (of each model) are fixed at α = 0.7, r = 0.7, K = 1000, τ = 2, and μ_0_ = 1000. Equal prior probabilities (1/6) are assigned to each FD model and we obtained 1,50,000 posterior samples for each model parameter. The convergence of all posterior distributions was assessed through the Gelman-Rubin convergence diagnostics^40^, by using the `gelman.diag()` function from `coda` ^40^ package in `R` software. Fig 2-B presents the posterior model probability of each FD model. Caputo FD (model *M*_1_) emerges as the most optimal FD with posterior model probability of *M*_1_ equals to 0.9993 for simulated data. Marginal densities of model parameters were derived from joint posterior distributions generated via JAGS and approximated with a Gaussian kernel density estimator. (Extended Fig 1).

We expanded the simulation experiments on simulated datasets using five distinct parameter configurations for each FD model (model *M*_1_ − *M*_6_). For each configuration, the prior distribution and initial values of the model parameters were maintained consistent with those specified for the Caputo FD model above. Only the simulation study corresponding to the data generated from model *M*_3_, the prior distribution assumptions for the parameters were made as *α* ∼ 𝒩(α_0_, (0.0001)^2^), *r* ∼ 𝒩(r_0_, (0.01)^2^), *K* ∼ 𝒩(K_0_, (100)^2^), σ ∼ Γ(0.01, 0.01), and *μ*_0_ ∼ 𝒰(10, 600), where *α*_0_, *r*_0_, and *K*_0_ are the parameter values used for the data generation using model *M*_3_. Extended Table 1 presents the detailed posterior model probabilities for each configuration, highlighting the robustness of the BaFOMS approach. The results demonstrate the method’s accuracy in identifying the FD definition initially employed in the simulated data generation.

Our approach successfully infers the most optimal FD model for all three ecological population and one epidemiological dataset. Fig 5-A displays the posterior model probabilities of each FD model within the BaFOMS framework. GL – FD (Model *M*_6_) emerged out as best model in all cases with posterior model probability **0.3898, 0.9736, 0.44** and **0.9044** respectively. Notably, for *Ursus americanus* and *Phalacrocorax carbo*, the RL FD (Model *M*_4_) also provides a strong fit to the data (Fig 3 and Fig 5-A).

**Figure 5.**
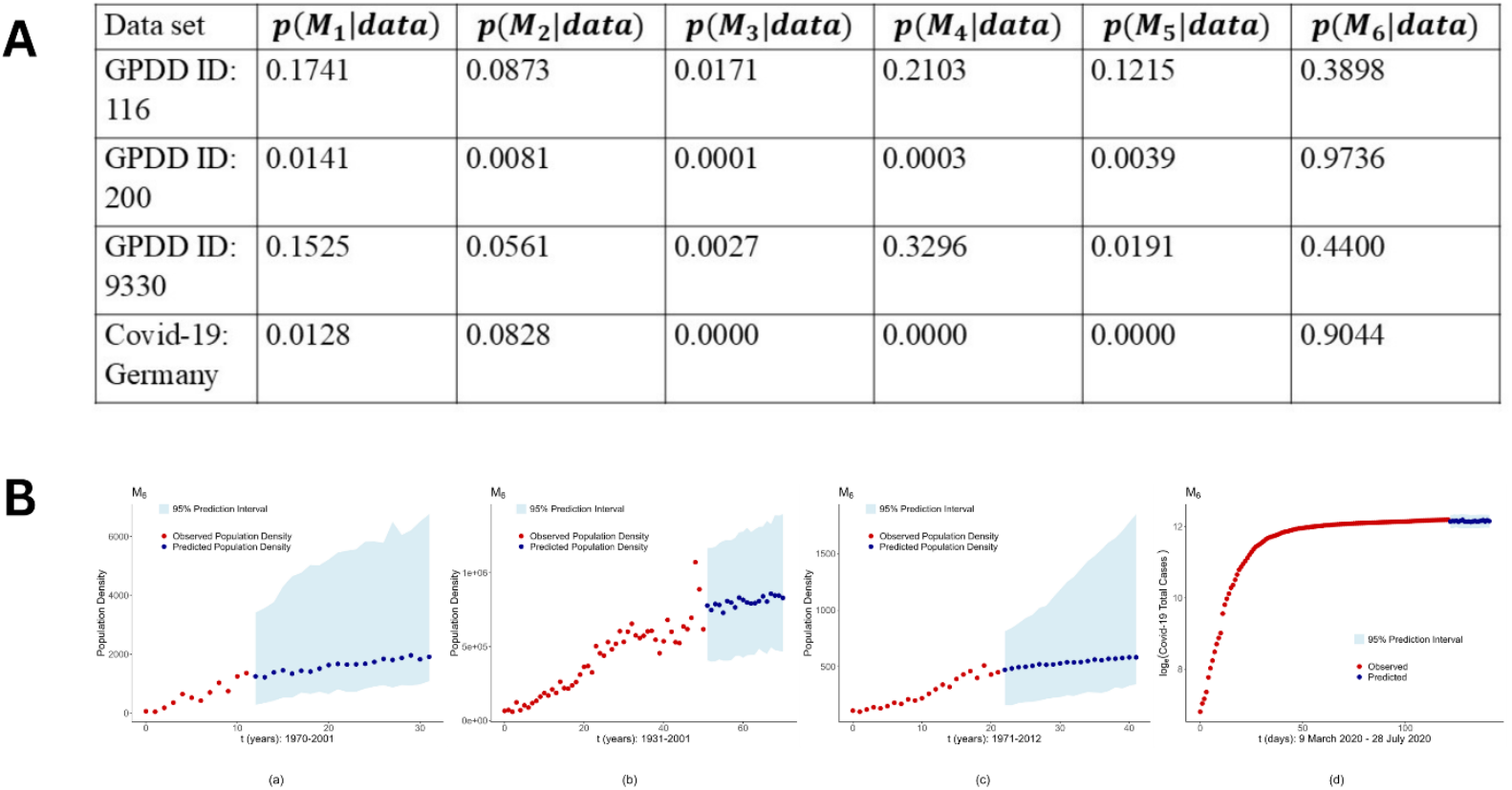
Optimal model section and forecasting for eco-epidemiological datasets. **A**. Posterior model probabilities were calculated using the RJMCMC method implemented with the rjmcmc package^35^ in R for four datasets: the ecological population datasets of Ursus americanus (GPDD ID: 116), Castor canadensis (GPDD ID: 200), Phalacrocorax carbo (GPDD ID: 9330), and the Covid-19 total cases dataset for Germany. **B**. Figures (a), (b), and (c) represent the species Ursus americanus (GPDD ID: 116), Castor canadensis (GPDD ID: 200), and Phalacrocorax carbo (GPDD ID: 9330), respectively. Figure (d) corresponds to the Covid-19 total cases data for Germany. Parameter values were sampled from the joint posterior distribution obtained using JAGS, and future values were projected over time. This process was repeated 1000 times, and the mode of the generated 1000 values at each time point was taken as the future predicted value. A 95% prediction interval was calculated using quantiles. Red solid points indicate the observed data, blue points show the predicted future values, and the light blue ribbon represents the 95% prediction interval.

### Forecasting with Uncertainty

We further examined the performance of BaFOMS on unseen datasets extending its functionality beyond model selection to include predictive analysis with quantifiable uncertainty. Using the best-fitted model (*M*_6_), we forecasted the evolution dynamic to subsequent 20 time points, accompanied by 95% prediction intervals for all datasets. Parameter values were sampled from the joint posterior distribution obtained via JAGS, ensuring that the inherent uncertainty in parameter estimation was fully accounted for during prediction. This process was repeated 1,000 times, with the 2.5th and 97.5th percentiles of the simulated population values used to derive the posterior predictive intervals (Fig 5-B). This predictive validation confirms that BaFOMS not only identifies the best-fitting model for historical data but also reliably forecast future trends.

Although, we restricted ourselves to predict short-term dynamics due to small training data and the underlying logistic growth model assumption, the overall architecture is adaptable and could be extended to forecast long-term evolutionary dynamics.

### Estimation of Parameters

Fractional models although effective in capturing continuous memory within dynamical systems, lacks a robust computation framework to solve the inverse problem of inferring model parameters from data ^16^. The complexity of the time correlation function and corresponding discretization scheme further complicates the estimation of parameters. One of the inherent advantages of the BaFOMS architecture is its ability to estimate robust parameter with well-defined credible intervals within the Bayesian framework. Model parameter estimation was performed using the posterior distribution obtained via JAGS^33^ for the most optimal model (*M*_6_) for all datasets. The mode of posterior distribution was taken as the point estimate of model parameters. To quantify the uncertainty associated with these estimates, the 95% posterior credible intervals were calculated by computing 2.5^th^ and 97.5^th^ percentiles of posterior samples (Fig 6-A). Fig 6-B also displays the posterior density function using kernel density approximation for all the parameters of most optimal model. Further we extended the analysis and reported the posterior density function and associated uncertainty for each model for every dataset (Supplementary/S3). We note that, although previous studies^31,41,42^ had focused on parameter estimation including the order α of the FD models using a Bayesian framework, BaFOMS extends these foundations by providing a comprehensive framework that addresses model selection, uncertainty, and parameter estimation in a unified manner.

**Figure 6.**
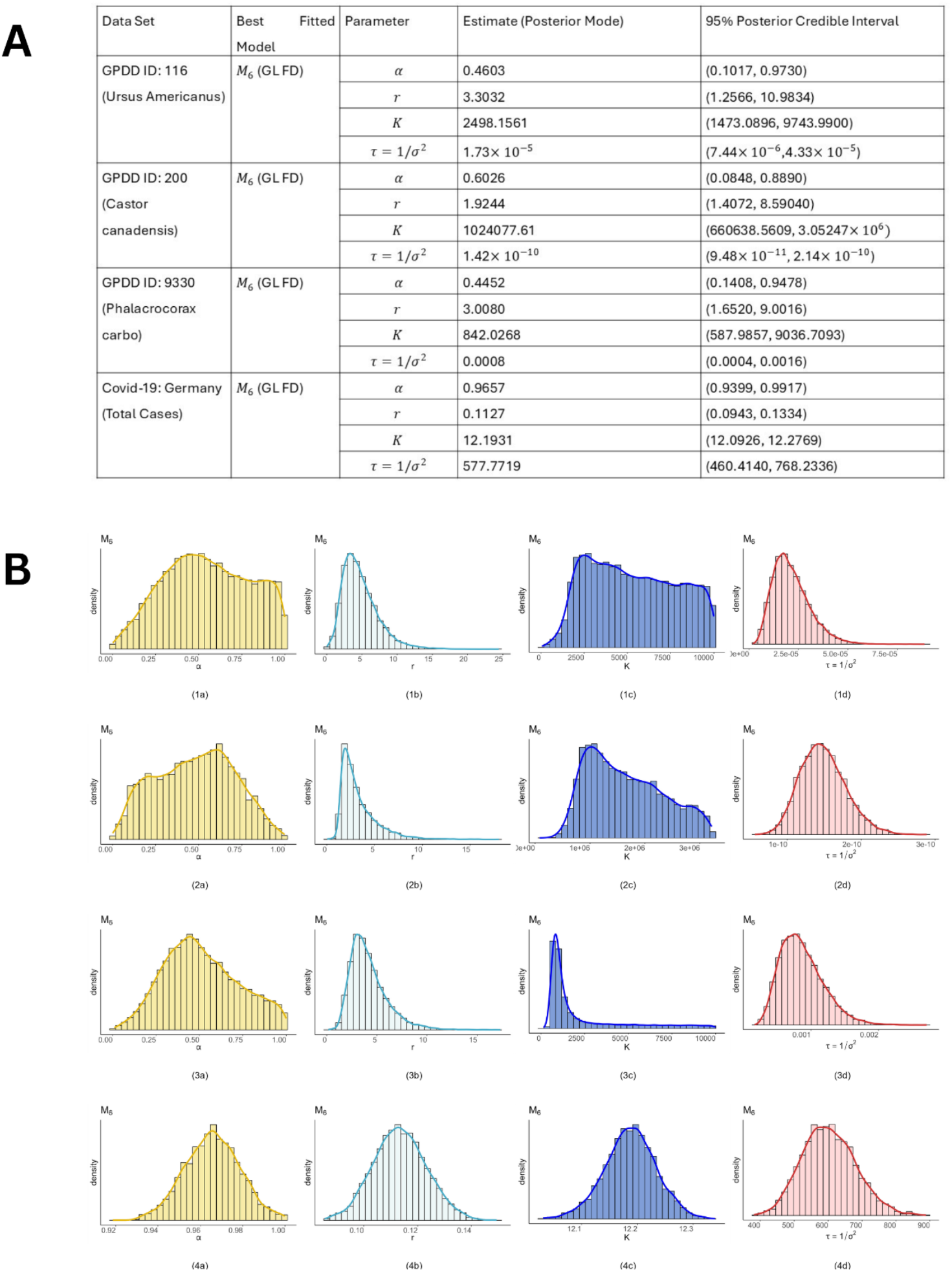
Posterior density of estimated parameters for best fitted model. **A**. Posterior density estimate of all parameters for all four eco-epidemiological datasets under the most optimal fitted model. These estimates were calculated as the mode of the posterior distribution derived through JAGS, with 95% posterior credible intervals computed from the 2.5th and 97.5th percentiles of the posterior distribution. **B**. Posterior Density plots for model parameters α, r, K, τ of best-fitted model (GL FD) for the studied datasets are shown. The first row (1a, 1b, 1c, and 1d) represents Ursus americanus, the second row (2a, 2b, 2c, and 2d) corresponds to Castor canadensis, the third row (3a, 3b, 3c, and 3d) depicts Phalacrocorax carbo, and the fourth row (4a, 4b, 4c, and 4d) represents the Covid-19 total cases data for Germany.

## Discussion

Many studies have explored aspects such as model uncertainty within a single FD definition^11,42^, hypothesis testing for selecting between FDE and ODE^31^, and parameter estimation, including the order (*α*) of FDE^41^. However, the uncertainty inherent in the choice of FD definition remains unexplored. Our method addresses this gap and extends beyond previous approaches, offering a comprehensive framework that unifies model selection, uncertainty quantification, and parameter estimation in a Bayesian framework. The potential application of BaFOMS extends well beyond eco-epidemiological dynamics, offering potential solutions for a variety of problems in engineering and physical sciences. It can be readily applied to infer the most optimal memory kernel in eco-epidemiological processed driven by multiphasic equations. Long-term dynamics, in particular, are typically captured by multiphasic growth equations, unlike the uniphasic logistic growth model we have restricted ourselves to in this study. These systems are constructed as linear combinations of uniphasic growth equations and characterized by multiple inflection points^42–47^. For instance, our approach can be applied to a wide range of stage-structured epidemic models^16^ with fractional orders, enabling the selection of the most optimal model that effectively captures the disease history, its evolution, and control dynamics. The BaFOMS framework offers a distinct computational advantage by not only integrating memory effects but also selecting the most appropriate method for incorporating them within the FDE system, thereby providing a simpler yet effective alternative to traditional methods^31^ and complex neural network models^16^.

It is noteworthy that in all eco-epidemiological case studies, GL – FD (Model *M*_6_) consistently emerged as the model with the highest posterior probability. The discrete summation nature of GL – FD naturally aligns it with discrete or irregular time series data in eco-epidemiological contexts. This property allows GL-FD to assign stronger weights to recent data points, enhancing its ability to capture local trends and short-term dependencies. In contrast, other models that rely on continuous derivatives tend to smooth out these recent changes, reducing their effectiveness in handling data with high variability or short-term fluctuations. Furthermore, eco-epidemiological systems often start from complex or unknown initial states, and the impact of these initial conditions on model outputs can make these continuous derivatives less reliable without precise initial data. The discreteness of GL – FD reduces its sensitivity to initial conditions and emphasizes recent data points, making it more robust when precise initial data is unavailable or uncertain. Additionally, the GL-FD model’s discretization scheme is computationally simpler and numerically stable, that perform efficiently in iterative procedures, such as MCMC or variational inference, especially when dealing with large datasets or high-dimensional models.

This discrete memory nature of these eco-epidemiological process also aligns with the results of integer order models being more robust the fractional order models in previous studies^16,31^. This leads to an important question: could an ODE or delay system better support the observed data? This prompts the hypothesis test *H*_0_: *α* = 1 (ODE) against *H*_1_: *α* < 1(FDE). The posterior distribution of *α* corresponding to the best FD definition, is informative to answer this question. If the posterior mode of *α* is close to 1, this would support the suitability of an ODE model for the population dynamics in question. We suggest that the decision to select the ODE or FDE should be considered by testing the null hypothesis *H*_0_: *α* = 1 against *H*_1_: *α* < 1 and reporting the p-values or by providing the evidence of *H*_1_ against *H*_0_ in Bayesian setting^31^. For systems governed by fixed delay dynamics, we can rely on the posterior model probabilities within the BaFOMS framework, where parameter space of delayed model will have no parameter as *α* characterizing the strength of memory. This will be a subject of our future studies.

Several promising directions for future research could further enhance the robustness and generalizability of the proposed method. In this study, we have kept the memory parameter *α*(*t*) constant to simplify computations. However, in many dynamical systems, memory effects evolve over time. A more suitable approach might involve a time-dependent fractional order model, enabling the estimation of a variable *α*(*t*). Previous studies, such as those by Jahanashi et al. ^48^ and Kharazmi et al. ^16^, have explored methods to estimate *α*(*t*), either by using a piecewise constant function with a few constant parameters or through complex physics-informed neural networks. These approaches, however, have limitations, including under-parameterization of fractional orders or highly complex computational schemes. The BaFOMS framework could address these challenges by offering a simpler and more efficient approach to estimate *α*(*t*). Another potential advancement is to tackle model uncertainty by considering multiple models under various FD definitions, which would yield more robust insights and predictive inference. For instance, if we seek to select the optimal growth model from a set (e.g., logistic, Gompertz, exponential, theta-logistic, Allee) ^11,49,50^, each model could be evaluated under six different FD definitions. This approach would produce a complete model space containing 30 models, allowing for inference based on the posterior distribution across this model space and addressing uncertainty both in model selection and FD definition.

## Methods

### Ecological population data

Yearly population count time series data for three species were obtained from the Global Population Dynamic Database (GPDD)^38^. GDPP contains 4471 time series on the population abundance of plant and animal species worldwide, covering a total of 1891 taxonomic groups, collected from 988 different locations. The data was accessed from the Global Population Dynamics Database (GPDD) as CSV files via the KNB repository^38^. We retrieved temporal sequences representing annual population density values for three distinct taxa. Specifically, two datasets pertain to mammalian species: `Ursus Americanus` (commonly known as the American black bear, GPDD Main ID: 116), spanning 12 years (1970-1981), and `Castor canadensis` (commonly known as the beaver, GPDD Main ID: 200), spanning 51 years (1931-1981). Concurrently, the avian dataset corresponds to `Phalacrocorax carbo` (commonly known as the great cormorant, GPDD Main ID: 9330), with a temporal extent of 21 years (1971-1992). For the dataset corresponding to the great cormorant (GPDD Main ID 9330), the population density count for the year 1973 was missing. Therefore, we imputed this value using the mean of the population density counts from the previous and next years and carried out the analysis. We have extracted the data for each species and stored it in separate csv files located at https://github.com/dipalimestry96/BaFOMS-Bayesian-Fractional-Order-Model-Selection. These files can be accessed using the guide provided in the supplementary material (Supplementary/S4).

### Covid-19 data

The Covid-19 case data^39^, spans from 31^*St*^ December 2019 to 8^*th*^ July 2020, encompassing records of confirmed Covid-19 cases during first outbreak in Germany (https://ourworldindata.org/coronavirus). Notably, up to 27^*th*^ January 2020, Germany had not reported any instances of Covid-19. Additionally, minimal fluctuations in the total confirmed cases were observed until 8^*th*^ March 2020, after which the data exhibited a sigmoidal trend consistent with our model assumption. Consequently, we confined our analysis to the period from 8^*th*^ March 2020, to 8^*th*^ July 2020. It’s noteworthy that all investigations conducted on this dataset employed a natural logarithmic scale for the confirmed total cases.

### Model development

#### Mathematical model

A simple fractional order initial value logistic model is considered as

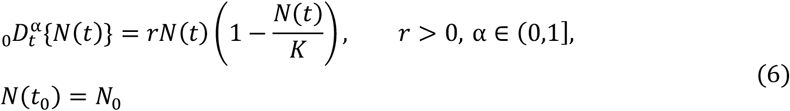

Where *N*(*t*) represents the population size at time t; *N*_0_ is the initial population size; *r* is the intrinsic rate of growth; *K* is the carrying capacity of the system; and 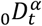 denotes one of the FDs listed in Figure 1-B. For α = 1, equation (6) reduces to well-known logistic growth equation whose solution is given by 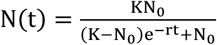. However, for 0 < α < 1, there is no closed-form analytical solution, requiring numerical approximation methods for solving the fractional logistic equation. Two-step Adams-Bashforth scheme^51^ is applied to discretize the FDs which are defined in Caputo sense such as Caputo, CF, and ABC. RL, CF-RL being defined in RL sense are discretized using forward difference followed by numerical approximation of integral using Trapezoidal rule^25^. Finite difference method using Taylor’s approximation^41^ is applied to numerically discretize the discrete nature of GL fractional logistic equation. Importantly, RL and GL fractional derivatives as distinct definitions in this context, allowing us to discern alterations in posterior distribution and convergence probability in Bayesian estimation due to the discrete nature of the GL definition. While Hernandez et al.^41^ briefly discussed Bayesian inference by numerically approximating the RL FD using the GL definition, our approach treats them as separate entities, facilitating a comprehensive exploration of nuances in Bayesian estimation. The complete discretization scheme along with fractional properties of these FDs are discussed in (Supplementary/S2.1, S2.2, S2.3).

### Bayesian Analysis

#### Bayesian Estimation

The posterior distribution is fundamental to Bayesian analysis, which involves estimating the probability distribution of unknown parameters (*θ*) based on observed data and prior knowledge. It is computed using Bayes’ theorem, which combines the data likelihood (the probability of observing the data given a particular set of parameters) and the prior distribution (representing the initial beliefs about parameter values before observing the data) as

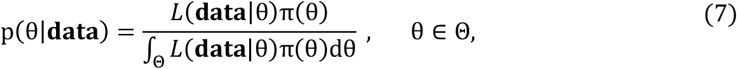

where *L*(**data**|*θ*) is the likelihood function and *π*(*θ*) is the prior distribution of *θ*. The integral over the parameter space θ ensures the normalization constant of the posterior distribution.

#### Likelihood function

The applied numerical scheme for approximating FDE solutions, resulted in distribution of the current state variable being dependent on previous time points. This dependency prevents expressing the data likelihood as a product of marginal likelihoods. Instead, leveraging the Markov property, the data likelihood is represented as a product of conditional likelihood functions, as described below:

##### Case 1

For the models (***M***_**1**_, ***M***_**2**_, ***M***_**3**_), the present size depends on the previous two state variables values. Therefore, the likelihood function for the dataset 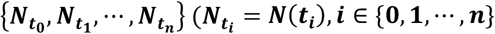 from the model given in equation (3) corresponding to the ***m*** = **1, 2, 3** is as follows:

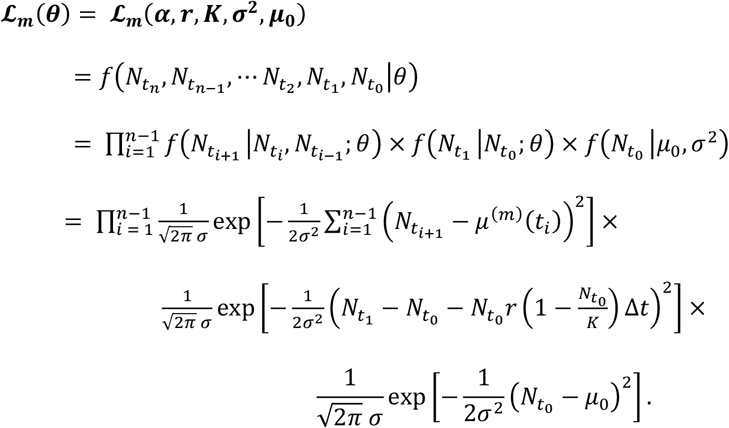

##### Case 2

The Markovian structure of the present state variable value for the last three models (***M***_**4**_ − ***M***_**6**_) depends on state variables values on all preceding time points (starting of the process). Therefore, the likelihood function corresponding to these models (***m*** = **4, 5, 6**) is as follows:

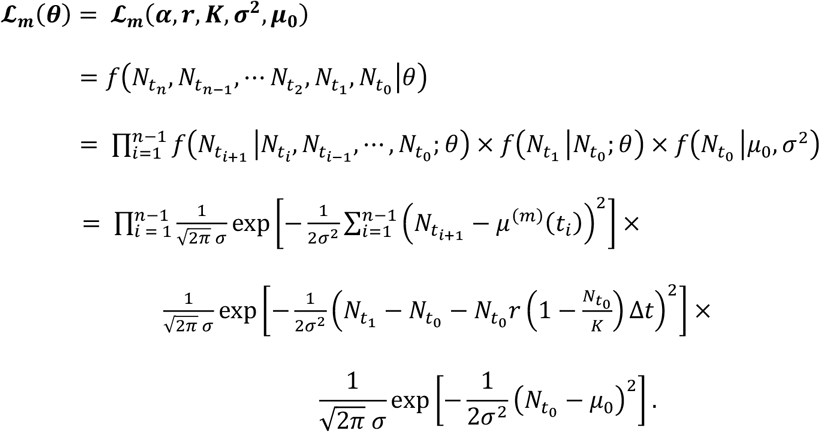

All six models (equation (3), for *m* ∈ {1,2, ⋯ 6}) are highly nonlinear, making parameter estimation and model selection challenging with traditional statistical techniques. Therefore, we employ Bayesian methods to address these complexities effectively.

## Data Availability

The ecological time series data is publicly available at Global Population Dynamic Database (GPDD)^38^. The empirical Covid-19 pandemic data for Germany^39^ can be accessed from https://ourworldindata.org/coronavirus. All simulated and experimental datasets are publicly available at https://github.com/dipalimestry96/BaFOMS-Bayesian-Fractional-Order-Model-Selection/tree/main/Data%20Analysis%20Mestry%20et%20al%202024. The simulated datasets were generated according to the approach described in “Results” section.

## Code Availability

The code for reproducing the analyses and figures in this paper, as well as for applying the methodology we have developed here, are freely available at https://github.com/dipalimestry96/BaFOMS-Bayesian-Fractional-Order-Model-Selection.

## Acknowledgments

Mestry D. V. is thankful to the DST-INSPIRE, Government of India, for the financial support in the form of a research fellowship (Application Ref. No. DST/INSPIRE/03/2019/000445, Inspire Reg. No. IF190606).

## Extended Figures and Table

**Extended Figure 1.**
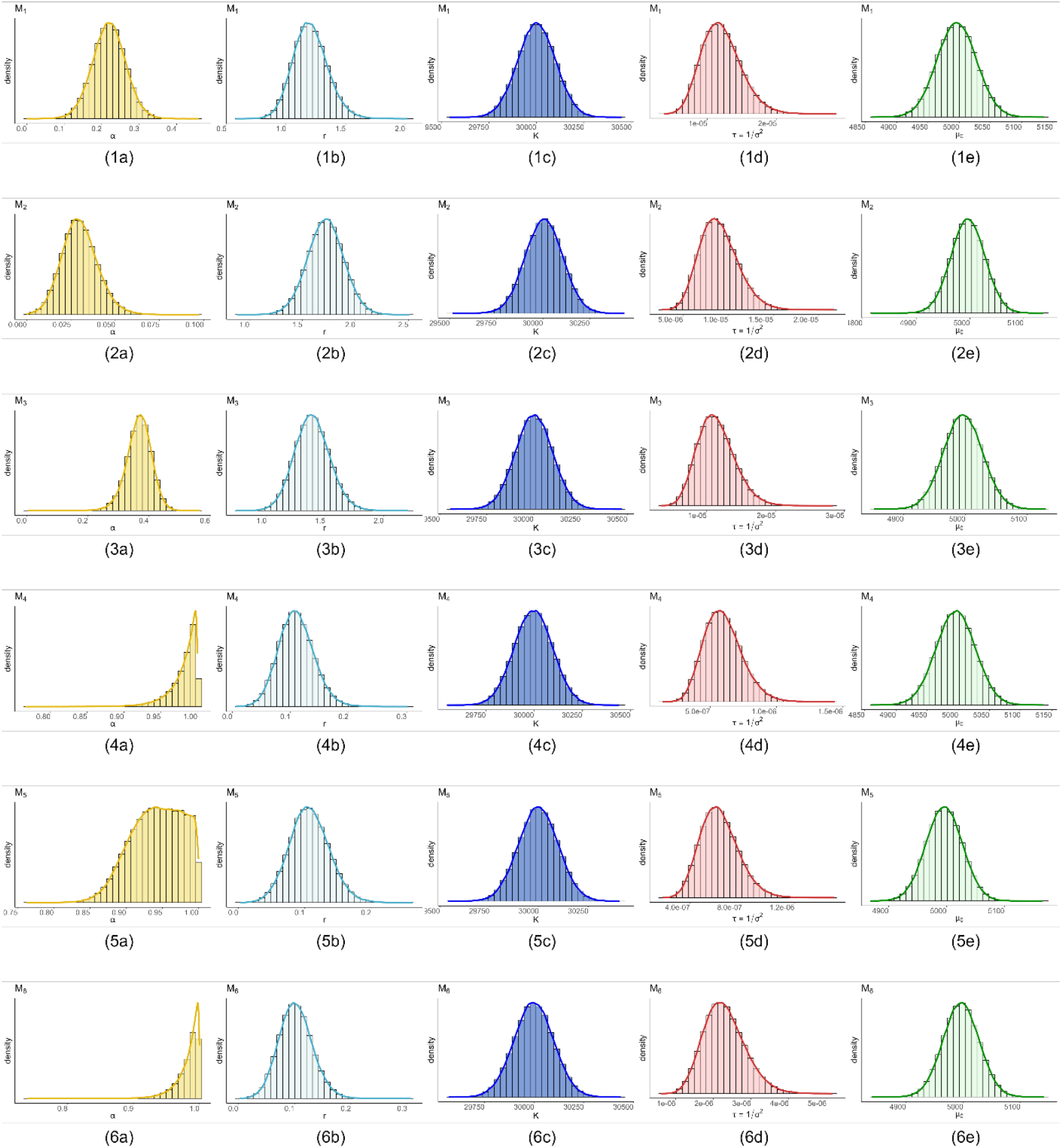
Posterior density plot of model’s parameters under simulated dataset using. Model M_1_ (Caputo FD). The rows one through six represent the posterior distributions of the model parameters (α, r, K, τ, μ_0_)′ for models M_1_ to M_6_. Each parameter subplot displays a histogram of the posterior distribution with an overlaid density curve, derived using a Gaussian kernel density estimator. The dataset comprising 40 observations was generated using the parameter settings (α, r, K, σ^2^, μ_0_)^′^ = (0.3, 1.1, 30000, 300^2^, 5000)′ over the interval t ∈ [0, 20] from model M_1_ (Caputo FD).

**Extended Table 1.**
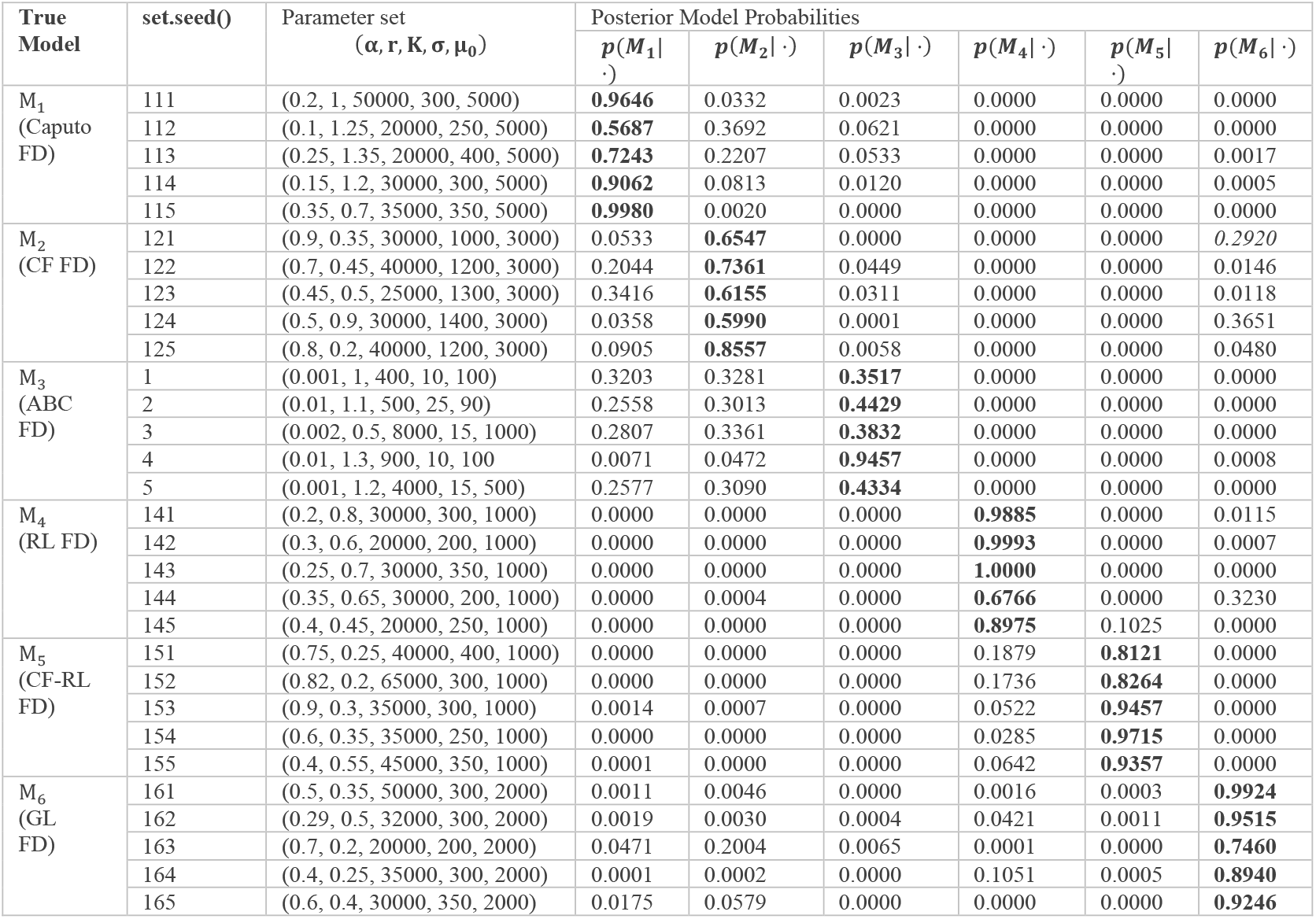
Posterior probability of all models from simulated data generated by each model. The table presents posterior probabilities obtained through the Bayesian model selection approach in a comprehensive simulation experiment. Five different parameter sets were used sequentially to generate data and conduct the simulation study. This process was repeated for all six models M_1_ − M_6_. The seed values, indicated in the second column, are reported to ensure the repeatability of the results shown in the table.

