## Supplementary Information for "Deciphering Memory Patterns in Eco-Epidemiological systems through the lens of BaFOMS: A Bayesian Fractional Order Model Selection Method"

---

---

**S3 Parameter Distributions of Alternative Models: Real Data Analysis** **15**

**S4 Fractional Differential definition selection (R codes)** **19**

---

[*]Corresponding author

[**#]These authors contributed equally to this work and share first authorship.

[1]Institute of Chemical Technology, Mumbai, Maharashtra, India

[2]Interdisciplinary Center for Scientific Computing, University of Heidelberg, Germany

[3]Heidelberg Institute of Global Health (HIGH), Universitätsklinikum Heidelberg, Heidelberg - 69120, Baden Württemberg, Germany.

### S1. Fractional Differential Equations : Introduction and Preliminaries

This section will systematically introduce some basic definitions of fractional derivatives, preliminary concepts, theorems, and a few basic assumptions. This will collectively serve as the framework upon which we will construct the mathematical concept of memory precisely.

**Definition S1.1.** *(Atangana and Owolabi, 2018; Caputo, 1967) The Caputo fractional derivative with order $\alpha$ of a function $\phi \in H^1(a,b)$, $b > a$, and $a \in (-\infty, t)$ is given as*

$$ {}_{a}^{C}D_t^{\alpha}\{\phi(t)\} = \frac{1}{\Gamma(n-\alpha)} \int_a^t \frac{1}{(t-k)^{\alpha-n+1}} \phi^{(n)}(k)\, dk, \tag{1}$$

*where $n-1 \leq \alpha \leq n \in \mathbb{N}$, $t \geq a$, and $H^1(a,b)$ is a Hilbert space.*

*If $\alpha \in [0,1]$ and $\phi \in H^1(0,b)$, $b > 0$ then*

$$ {}_{0}^{C}D_t^{\alpha}\{\phi(t)\} = \frac{1}{\Gamma(1-\alpha)} \int_0^t \frac{1}{(t-k)^{\alpha}} \phi^{'}(k)\, dk, \tag{2}$$

*where $t \geq 0$.*

**Definition S1.2.** *(Atangana and Gómez-Aguilar, 2018; Podlubny, 1999) The R-L derivative of a function $\phi(t)$ (not necessarily differentiable) with fractional order $n-1 < \alpha \leq n \in \mathbb{Z}^+$ is defined as*

$$ {}_{0}^{RL}D_t^{\alpha}\{\phi(t)\} = \frac{1}{\Gamma(n-\alpha)} \frac{d^n}{dt^n} \int_0^t \phi(k)(t-k)^{n-\alpha-1}\, dk. \tag{3}$$

*If $\alpha \in (0,1]$, then*

$$ {}_{0}^{RL}D_t^{\alpha}\{\phi(t)\} = \frac{1}{\Gamma(1-\alpha)} \frac{d}{dt} \int_0^t \phi(k)(t-k)^{-\alpha}\, dk. \tag{4}$$

Consider a Fractional Initial Value Problem (FIVP) in the Caputo sense as

$$ {}_{0}^{C}D_t^{\alpha}\{\phi(t)\} = f(t,\phi(t)), \quad \phi^m(0) = \phi_0^m, \tag{5}$$

where $m = 0, 1, 2, \cdots, \lceil\alpha\rceil - 1$ and $\lceil\alpha\rceil$ is the smallest integer $\geq \alpha$.
The FIVP in equation (5) can be rewritten in terms of Volterra integral equation (Diethelm and Ford, 2004) as

$$ \phi(t) = \sum_{m=0}^{\lceil\alpha\rceil-1} \frac{t^m}{m!}\phi^m(0) + \frac{1}{\Gamma(\alpha)} \int_0^t [(t-k)^{\alpha-1} f(k,\phi(k))]\, dk. \tag{6}$$

If $0 \leq \alpha \leq 1$, then it is obvious to generalize the method for the classical first-order differential equation to FIVP appropriately. Therefore, extending equation (6) (Batogna and Atangana, 2017) as

$$ \phi(t) = \phi_0 + \frac{1}{\Gamma(\alpha)} \int_0^t [(t-k)^{\alpha-1} f(k,\phi(k))]\, dk. \tag{7}$$

**Definition S1.3.** *(Podlubny, 1999; Diethelm and Ford, 2004) The R-L integral operator of a function $\phi(t)$ with fractional order $\alpha > 0$ is defined as*

$$ {}_{0}^{RL}I_t^{\alpha}\{\phi(t)\} = \frac{1}{\Gamma(\alpha)} \int_0^t \phi(k)(t-k)^{\alpha-1}\, dk. \tag{8}$$

**Corollary S1.1.** *The Caputo (1) and R-L (3) fractional order derivative can also be expressed in terms of R-L integral operator as*

$$ {}_{a}^{C}D_{t}^{\alpha}\{\phi(t)\} \;=\; {}_{a}^{RL}I_{t}^{n-\alpha}D^{n}\{\phi(t)\}, \tag{9}$$

*and*

$$ {}_{0}^{RL}D_{t}^{\alpha}\{\phi(t)\} \;=\; D^{n}[{}_{0}^{RL}I_{t}^{n-\alpha}\{\phi(t)\}], \tag{10}$$

*where* $D^{n}=\frac{d^{n}}{dt^{n}}$ *is* $n^{th}$ *order differential operator.*

**Corollary S1.2.** *The Caputo (1) and R-L (3) fractional order derivative with order* $n-1\le\alpha\le n$ *are equivalent as*

$$ {}_{a}^{RL}D_{t}^{\alpha}\{\phi(t)\} \;=\; {}_{0}^{C}D_{t}^{\alpha}\{\phi(t)\} + \sum_{i=0}^{n-1}\frac{(t-b)^{(i-\alpha)}\phi^{(i)}(a)}{\Gamma(i-\alpha+1)}. \tag{11}$$

*The relation* ${}_{a}^{RL}D_{t}^{\alpha}\{\phi(t)\} \;=\; {}_{0}^{C}D_{t}^{\alpha}\{\phi(t)\}$ *holds only if initial condition* $\phi^{(i)}(a)=0,\; i=0,1,2,\ldots,n-1.$

**Definition S1.4.** *(Atangana and Owolabi, 2018; Caputo and Fabrizio, 2015) The Caputo-Fabrizio derivative of order* $\alpha$ *of a function* $\phi\in H^{1}(b_1,b_2)$, $b_2>b_1$, *and* $b_1\in(-\infty,t)$ *in caputo sense is defined as*

$$ {}_{b_1}^{CF}D_{t}^{\alpha}\{\phi(t)\} \;=\; \frac{\mathbb{M}(\alpha)}{(n-\alpha)}\int_{b_1}^{t}\phi^{(n)}(k)\,\exp\left(\frac{-\alpha(t-k)}{(n-\alpha)}\right)\,dk, \tag{12}$$

*where* $n-1\le\alpha\le n\in\mathbb{N}$, *and* $t\ge b_1$, $H^{1}(b_1,b_2)$ *is a Hilbert space and* $\mathbb{M}(\alpha)$ *is normalising function such that* $\mathbb{M}(0)=\mathbb{M}(1)=1.$

*If* $\alpha\in[0,1]$ *and* $\phi\in H^{1}(0,b)$, $b>0$ *then*

$$ {}_{0}^{CF}D_{t}^{\alpha}\{\phi(t)\} \;=\; \frac{\mathbb{M}(\alpha)}{(1-\alpha)}\int_{0}^{t}\phi^{'}(k)\,\exp\left(\frac{-\alpha(t-k)}{(1-\alpha)}\right)\,dk, \tag{13}$$

*where* $t\ge0.$

**Definition S1.5.** *(Atangana and Owolabi, 2018; Caputo and Fabrizio, 2015) The Caputo-Fabrizio derivative of order* $\alpha$ *of a function* $\phi\notin H^{1}(b_1,b_2)$, $b_2>b_1$, $b_1\in(-\infty,t)$ *and* $0\le\alpha\le1$ *in Caputo sense is defined as*

$$ {}_{b_1}^{CF}D_{t}^{\alpha}\{\phi(t)\} \;=\; \frac{\mathbb{M}(\alpha)}{(1-\alpha)}\int_{b_1}^{t}[\phi(t)-\phi(k)]\,\exp\left(\frac{-\alpha(t-k)}{(1-\alpha)}\right)\,dk, \tag{14}$$

*where* $H^{1}(b_1,b_2)$ *is a Hilbert space and* $\mathbb{M}(\alpha)$ *is normalising function such that* $\mathbb{M}(0)=\mathbb{M}(1)=1.$

Consider a Caputo - Fabrizio FIVP in the Caputo sense with order $\alpha\in[0,1]$, $t\ge0$ as

$$ {}_{0}^{CF}D_{t}^{\alpha}\{\phi(t)\} \;=\; f(t,\phi(t)). \tag{15}$$

Rewriting (15) using definition (S1.4) as

$$ \frac{\mathbb{M}(\alpha)}{(1-\alpha)}\int_{0}^{t}\phi^{'}(k)\,\exp\left(\frac{-\alpha(t-k)}{(1-\alpha)}\right)\,dk \;=\; f(t,\phi(t)), \tag{16}$$

or equivalently

$$\int_0^t \phi^{'}(k) \exp\left(\frac{\alpha}{(1-\alpha)}k\right) dk \;=\; \exp\left(\frac{\alpha}{(1-\alpha)}t\right) f(t,\phi(t)). \tag{17}$$

By differentiating both sides and rearranging, we get

$$\phi^{'}(t) \;=\; \frac{1-\alpha}{\mathbb{M}(\alpha)} f^{'}(t,\phi(t)) + \frac{\alpha}{\mathbb{M}(\alpha)} f(t,\phi(t)). \tag{18}$$

Again integrating from 0 to $t$, we get

$$\phi(t) \;=\; \phi(0) + \frac{1-\alpha}{\mathbb{M}(\alpha)}[f(t,\phi(t)) - f(0,\phi(0))] + \frac{\alpha}{\mathbb{M}(\alpha)} \int_0^t f(k,\phi(k))\, dk. \tag{19}$$

**Definition S1.6.** *(Caputo and Fabrizio, 2015) The corresponding CF fractional integral in Caputo sense with order $0 \le \alpha \le 1$, $t \ge 0$ is given as*

$$^{CF}_{\;\,0}I^{\alpha}_{t}\{\phi(t)\} \;=\; \frac{1-\alpha}{\mathbb{M}(\alpha)}\phi(t) + \frac{\alpha}{\mathbb{M}(\alpha)} \int_0^t \phi(k)\, dk. \tag{20}$$

**Definition S1.7.** *(Atangana and Gómez-Aguilar, 2018; Li et al., 2011) The CF (exponential decay) fractional derivative with order $n-1 \le \alpha \le n$, $t \ge 0$ of a function $\phi(t)$ (not necessarily differentiable) in R-L sense is given as*

$$^{CFR}_{\;\;\;\,0}D^{\alpha}_{t}\{\phi(t)\} \;=\; \frac{\mathbb{M}(\alpha)}{(n-\alpha)} \frac{d^n}{dt^n} \int_0^t \phi(k)\, \exp\left(\frac{-\alpha(t-k)}{(n-\alpha)}\right) dk, \tag{21}$$

*where $\mathbb{M}(\alpha)$ is normalising function such that $\mathbb{M}(0) = \mathbb{M}(1) = 1$.*

*If $\alpha \in [0,1]$, then*

$$^{CFR}_{\;\;\;\,0}D^{\alpha}_{t}\{\phi(t)\} \;=\; \frac{\mathbb{M}(\alpha)}{(1-\alpha)} \frac{d}{dt} \int_0^t \phi(k)\, \exp\left(\frac{-\alpha(t-k)}{(1-\alpha)}\right) dk. \tag{22}$$

**Remark S1.** *Fractional order modeling problems often require integration over the order of the fractional order system* i.e *$\alpha$, and it is desirable to have some values of $\alpha$ to contribute more than others. Hence, the multiplier function $\mathbb{M}(\alpha)$ is introduced such that $\mathbb{M}(0) = \mathbb{M}(1) = 1$. $\mathbb{M}(\alpha)$ could be any normalisation function satisfying $\mathbb{M}(0) = \mathbb{M}(1) = 1$, but in present work $\mathbb{M}(\alpha) = 1$ will be a constant function for simplicity (Baleanu and Fernandez, 2017).*

**Definition S1.8.** *(Atangana and Owolabi, 2018; Atangana, 2016) The Atangana-Baleanu (AB) fractional derivative of order $0 \le \alpha \le 1$ of a function $\phi \in H^1(b_1, b_2)$, $b_2 > b_1$, and $b_1 \in (-\infty, t)$ in Caputo sense is defined as*

$$^{ABC}_{\;\;\;b_1}D^{\alpha}_{t}\{\phi(t)\} \;=\; \frac{ABC(\alpha)}{1-\alpha} \int_{b_1}^t \phi^{'}(k)\, E_{\alpha}\left[\frac{-\alpha(t-k)^{\alpha}}{1-\alpha}\right] dk, \tag{23}$$

*where $E_\alpha$ is the generalised Mittag - Leffler function is given as $E_\alpha(-t^\alpha) = \sum_{m=0}^{\infty} \frac{(-t)^{\alpha m}}{\Gamma(\alpha m+1)}$ and $ABC(\alpha)$ is a normalising function having same properties and values as $M(\alpha)$ in remark (S1).*

**Definition S1.9.** *(Atangana, 2016) The corresponding Atangana-Baleanu (AB) fractional integral of order $0 \le \alpha \le 1$ with on local kernel of a function $\phi$ in Caputo sense is defined as*

$$^{ABC}_{\;\;\;\,0}I^{\alpha}_{t}\{\phi(t)\} \;=\; \frac{1-\alpha}{ABC(\alpha)}\phi(t) + \frac{\alpha}{ABC(\alpha)\Gamma(\alpha)} \int_0^t (t-k)^{(\alpha-1)}\phi(k)\, dk, \tag{24}$$

*where $ABC(\alpha)$ is a normalising function having same properties and values as $M(\alpha)$ in remark (S1).*

**Definition S1.10.** *(Podlubny, 1999; Li et al., 2011) The G-L fractional derivative of order $n-1 \leq \alpha \leq n$ of a function $\phi$ is given as*

$$\begin{aligned} {}_a^{GL}D_t^{\alpha}\{\phi(t)\} &= \lim_{\substack{h \to 0 \\ mh=t}} h^{-\alpha} \sum_{k=0}^{m} (-1)^k \binom{\alpha}{k} \alpha(t-kh), \quad m = 0, 1, 2, \cdots \\ &= \sum_{r=0}^{n-1} \frac{\phi^{(r)}(a)(t-a)^{(r-\alpha)}}{\Gamma(r+1-\alpha)} + \frac{1}{\Gamma(n-\alpha)} \int_0^t \frac{1}{(t-a)^{\alpha-n+1}} \phi^{(n)}(a)\, da, \end{aligned} \tag{25}$$

*where* $\binom{\alpha}{k} = \frac{\Gamma(\alpha+1)}{\Gamma(k+1)\Gamma(\alpha-k+1)}$.

**Corollary S1.3.** *It follows from the equation (25) and corollary (S1.2) that R-L and G-L derivatives are equivalent. In fact, equation (25) is a numerical method to solve R-L fractional derivative problems (21).*

**Remark S2.** *The present text will treat R-L (definition S1.7) and G-L (definition S1.10) fractional derivatives as two separate definitions only to apprehend changes in posterior and convergence probability in Bayesian estimation due to the discrete nature of G-L definition. Ariza-Hernandez et al. (2020a) provided a brief discussion from the point of view of the Bayesian inference where the R-L fractional derivative (S1.2) is numerically approximated using the G-L (S1.10) definition.*

### S2. Numerical discretization

Unlike ordinary derivatives, fractional derivatives are characterized by their hereditary nature, representing functions that maintain a comprehensive memory of prior states. Due to its complex nature, it becomes very difficult to solve FDEs analytically and sometimes analytical solution is not even feasible. Consequently, a substantial research has been put in exploration of approximate solutions for FDEs through the utilization of numerical methods both space (Owolabi, 2016; Zeng, 2015) and in time (Baleanu et al., 2016; Atangana, 2017). This section will explore the approximate numerical solution for FDEs with different type of Fractional operator in order to compute corresponding likelihood functions for fractional order logistic model (reference to main paper)

#### *S2.1. Caputo type FDEs*

FDEs with fractional operator defined in caputo sense, specifically Caputo (S1.1), C-F (S1.4), A-B (S1.8) are discretized numerically using the well established Adams - Bashforth scheme (Li and Tao, 2009). Atangana and Owolabi (2018) first explored the numerical solution of Caputo type FDEs followed by a corrigendum (Atangana and Owolabi, 2021) to correct their previous published results. However, some disparities in revised numerical results were still noted in our analysis. Consequently, we opted to re-derive the numerically solution using Adams - Bashforth scheme following the approach outlined by Atangana and Owolabi (2018). To further validate the accuracy of numerical solutions, we conducted an an additional verification step by substituting the order $\alpha$ of fractional derivative equal to 1, effectively recovering the classical Adams-Bashforth scheme. The convergence and stability results for the Adams - Bashforth scheme are discussed in Atangana and Owolabi (2018).

##### *S2.1.1. Caputo fractional derivative*

Consider the FDE of the form

$$\begin{aligned} {}_0^{C}D_t^{\alpha} y(t) &= f(t, y(t)), \;\; \alpha \in (0, 1], \\ y(0) &= y_0. \end{aligned} \tag{26}$$

Applying the generalized method of solution of FIVP (eqn. 6) on eqn. (26), we get

$$y(t) - y(0) \;=\; \frac{1}{\Gamma(\alpha)} \int_0^t [(t-k)^{\alpha-1} f(k, y(k))]\, dk, \tag{27}$$

So, at $t = t_{n+1}$, where $n = 0, 1, 2, ...,$ we obtain

$$y(t_{n+1}) - y(0) \;=\; \frac{1}{\Gamma(\alpha)} \int_0^{t_{n+1}} [(t_{n+1}-t)^{\alpha-1} f(t, y(t))]\, dt \tag{28}$$

and at $t = t_n$

$$y(t_n) - y(0) \;=\; \frac{1}{\Gamma(\alpha)} \int_0^{t_n} [(t_n-t)^{\alpha-1} f(t, y(t))]\, dt \tag{29}$$

Subtracting eqn. (29) from eqn. (28) yields

$$y(t_{n+1}) \;=\; y(t_n) + \frac{1}{\Gamma(\alpha)} \int_0^{t_{n+1}} [(t_{n+1}-t)^{\alpha-1} f(t, y(t))]\, dt \;-\; \frac{1}{\Gamma(\alpha)} \int_0^{t_n} [(t_n-t)^{\alpha-1} f(t, y(t))]\, dt. \tag{30}$$

For simplicity, we will use the form

$$y(t_{n+1}) \;=\; y(t_n) + A_1 - A_2 \tag{31}$$

where

$$A_1 \;=\; \frac{1}{\Gamma(\alpha)} \int_0^{t_{n+1}} [(t_{n+1}-t)^{\alpha-1} f(t, y(t))]\, dt \tag{32}$$

and

$$A_2 \;=\; \frac{1}{\Gamma(\alpha)} \int_0^{t_n} [(t_n-t)^{\alpha-1} f(t, y(t))]\, dt, \tag{33}$$

Upon approximating the $f(t, y(t))$ using the Lagrange interpolation and ignoring the approximation error, we an write $A_1$ as

$$\begin{aligned} A_1 \;&=\; \frac{1}{\Gamma(\alpha)} \int_0^{t_{n+1}} (t_{n+1}-t)^{\alpha-1} \left[ \frac{f(t_n, y_n)}{h}(t - t_{n-1}) \;-\; \frac{f(t_{n-1}, y_{n-1})}{h}(t - t_n) \right] dt \\ &=\; \frac{f(t_n, y_n)}{h\Gamma(\alpha)} \int_0^{t_{n+1}} (t_{n+1}-t)^{\alpha-1}(t - t_{n-1})dt - \frac{f(t_{n-1}, y_{n-1})}{h\Gamma(\alpha)} \int_0^{t_{n+1}} (t_{n+1}-t)^{\alpha-1}(t - t_n)dt \\ &=\; \frac{f(t_n, y_n)}{h\Gamma(\alpha)} A_{11} - \frac{f(t_{n-1}, y_{n-1})}{h\Gamma(\alpha)} A_{12} \end{aligned} \tag{34}$$

where $h = t_n - t_{n-1}$ is the step size and,

$$
\begin{aligned}
A_{11} &= \int_0^{t_{n+1}} (t_{n+1} - t)^{\alpha-1}(t - t_{n-1})dt, \\
A_{12} &= \int_0^{t_{n+1}} (t_{n+1} - t)^{\alpha-1}(t - t_n)dt.
\end{aligned} \tag{35}
$$

Substituting $y = t_{n+1} - t$ in eqn. (35) and simplifying, we get $A_{11}$ and $A_{12}$ as

$$
\begin{aligned}
A_{11} &= \int_0^{t_{n+1}} y^{\alpha-1}(t_{n+1} - y)dy - \int_0^{t_{n+1}} y^{\alpha-1} t_{n-1} dy, \\
A_{12} &= \int_0^{t_{n+1}} y^{\alpha-1}(t_{n+1} - y)dy - \int_0^{t_{n+1}} y^{\alpha-1} t_n dy.
\end{aligned} \tag{36}
$$

Upon simplifying eqn. (36), it gives

$$
\begin{aligned}
A_{11} &= \frac{t_{n+1}^{\alpha+1}}{\alpha(\alpha+1)} - \frac{t_{n+1}^{\alpha} t_{n-1}}{\alpha}, \\
A_{12} &= \frac{t_{n+1}^{\alpha+1}}{\alpha(\alpha+1)} - \frac{t_{n+1}^{\alpha} t_n}{\alpha}.
\end{aligned} \tag{37}
$$

Thus, substituting $A_{11}$ and $A_{11}$ in eqn.(34), we get $A_1$ as

$$
A_1 = \frac{f(t_n, y_n)}{h\Gamma(\alpha)} \left[ \frac{t_{n+1}^{\alpha+1}}{\alpha(\alpha+1)} - \frac{t_{n+1}^{\alpha} t_{n-1}}{\alpha} \right] - \frac{f(t_{n-1}, y_{n-1})}{h\Gamma(\alpha)} \left[ \frac{t_{n+1}^{\alpha+1}}{\alpha(\alpha+1)} - \frac{t_{n+1}^{\alpha} t_n}{\alpha} \right] \tag{38}
$$

Similarly, upon approximating the $f(t, y(t))$ using the Lagrange interpolation as in $A_1$ (34) and ignoring the approximation error, we obtain $A_2$ as

$$
\begin{aligned}
A_2 &= \frac{1}{\Gamma(\alpha)} \int_0^{t_n} (t_n - t)^{\alpha-1} \left[ \frac{f(t_n, y_n)}{h}(t - t_{n-1}) - \frac{f(t_{n-1}, y_{n-1})}{h}(t - t_n) \right] dt \\
&= \frac{f(t_n, y_n)}{h\Gamma(\alpha)} \int_0^{t_n} (t_n - t)^{\alpha-1}(t - t_{n-1})dt - \frac{f(t_{n-1}, y_{n-1})}{h\Gamma(\alpha)} \int_0^{t_n} (t_n - t)^{\alpha-1}(t - t_n)dt \\
&= \frac{f(t_n, y_n)}{h\Gamma(\alpha)} A_{21} - \frac{f(t_{n-1}, y_{n-1})}{h\Gamma(\alpha)} A_{22}
\end{aligned} \tag{39}
$$

where

$$
\begin{aligned}
A_{21} &= \int_0^{t_n} (t_n - t)^{\alpha-1}(t - t_{n-1})dt, \\
A_{22} &= \int_0^{t_n} (t_n - t)^{\alpha-1}(t - t_n)dt.
\end{aligned} \tag{40}
$$

Substituting $y = t_{n+1} - t$ in eqn. (40) and upon simplifying, we get $A_{21}$ and $A_{22}$ as

$$A_{21} = \frac{t_n^{\alpha+1}}{\alpha(\alpha+1)} - \frac{t_n^{\alpha} t_{n-1}}{\alpha}, \tag{41}$$
$$A_{22} = \frac{t_n^{\alpha+1}}{(\alpha+1)}.$$

Therefore, substituting $A_{21}$ and $A_{21}$ in eqn.(39), we get $A_2$ as

$$A_2 = \frac{f(t_n, y_n)}{h\Gamma(\alpha)} \left[ \frac{t_n^{\alpha+1}}{\alpha(\alpha+1)} - \frac{t_n^{\alpha} t_{n-1}}{\alpha} \right] - \frac{f(t_{n-1}, y_{n-1})}{h\Gamma(\alpha)} \left[ \frac{t_n^{\alpha+1}}{(\alpha+1)} \right] \tag{42}$$

Thus the approximate solution can be evaluated by substituting $A_1$ and $A_2$ in (31) as

$$\begin{aligned} y(t_{n+1}) &= y(t_n) + \frac{f(t_n, y_n)}{h\Gamma(\alpha)} \left[ \frac{t_{n+1}^{\alpha+1}}{\alpha(\alpha+1)} - \frac{t_{n+1}^{\alpha} t_{n-1}}{\alpha} - \frac{t_n^{\alpha+1}}{\alpha(\alpha+1)} + \frac{t_n^{\alpha} t_{n-1}}{\alpha} \right] \\ &\quad - \frac{f(t_{n-1}, y_{n-1})}{h\Gamma(\alpha)} \left[ \frac{t_{n+1}^{\alpha+1}}{\alpha(\alpha+1)} - \frac{t_{n+1}^{\alpha} t_n}{\alpha} + \frac{t_n^{\alpha+1}}{\alpha+1} \right] \end{aligned} \tag{43}$$

**Remark S3.** *when $\alpha = 1$, eqn (43) gives*

$$\begin{aligned} y(t_{n+1}) &= y(t_n) + \frac{f(t_n, y_n)}{h} \left[ \frac{t_{n+1}^2}{2} - t_{n+1} t_{n-1} - \frac{t_n^2}{2} + t_n t_{n-1} \right] \\ &\quad - \frac{f(t_{n-1}, y_{n-1})}{h} \left[ \frac{t_{n+1}^2}{2} - t_{n+1} t_n + \frac{t_n^2}{2} \right] \end{aligned} \tag{44}$$

*Substituting $h = t_n - t_{n-1} = t_{n+1} - t_n$ and simplifying further, we recover the classical Adams-Bashforth scheme as*

$$y(t_{n+1}) = y(t_n) + \frac{3}{2} h f(t_n, y_n) - \frac{1}{2} h f(t_{n-1}, y_{n-1}) \tag{45}$$

**Remark S4.** *The approximate numerical solution derived in equation (43) through the Adams-Bashforth scheme exhibits notable disparities when compared to the initially solved (eqn 3.17) in Atangana and Owolabi (2018) and in revised solution (eqn (1.1) of Atangana and Owolabi (2021)).*

*S2.1.2. Caputo - Fabrizio fractional derivative*

Consider the general FDE with exponentially decaying memory as

$$\begin{aligned} {}_0^{CF}D_t^{\alpha} y(t) &= f(t, y(t)), \ \alpha \in (0, 1], \\ y(0) &= y_0. \end{aligned} \tag{46}$$

Using fundamental theorem of calculus and applying generalized method of solution (eqn. 19) on eqn (46), we get

$$y(t) - y(0) = \frac{1-\alpha}{\mathbb{M}(\alpha)} [f(t, y(t)) - f(0, y(0))] + \frac{\alpha}{\mathbb{M}(\alpha)} \int_0^t f(k, y(k))\, dk. \tag{47}$$

So, at $t = t_{n+1}$, where $n = 0, 1, 2, ...$, we obtain

$$y(t_{n+1}) - y(0) = \frac{1-\alpha}{\mathbb{M}(\alpha)}[f(t_n, y(t_n)) - f(0, y(0))] + \frac{\alpha}{\mathbb{M}(\alpha)} \int_0^{t_{n+1}} f(t, y(t))\, dt \tag{48}$$

and at $t = t_n$

$$y(t_n) - y(0) = \frac{1-\alpha}{\mathbb{M}(\alpha)}[f(t_{n-1}, y(t_{n-1})) - f(0, y(0))] + \frac{\alpha}{\mathbb{M}(\alpha)} \int_0^{t_n} f(t, y(t))\, dt \tag{49}$$

Subtracting eqn. (49) from eqn. (48) yields

$$y(t_{n+1}) = y(t_n) + \frac{1-\alpha}{\mathbb{M}(\alpha)}[f(t_n, y(t_n)) - f(t_{n-1}, y(t_{n-1}))] + \frac{\alpha}{\mathbb{M}(\alpha)} \int_{t_n}^{t_{n+1}} f(t, y(t))\, dt. \tag{50}$$

Upon approximating the $f(t, y(t))$ using the Lagrange interpolation and ignoring the approximated error, we an write

$$\begin{aligned} y(t_{n+1}) &= y(t_n) + \frac{1-\alpha}{\mathbb{M}(\alpha)}[f(t_n, y(t_n)) - f(t_{n-1}, y(t_{n-1}))] \\ &+ \frac{\alpha}{\mathbb{M}(\alpha)} \int_{t_n}^{t_{n+1}} \left[\frac{f(t_n, y_n)}{h}(t - t_{n-1}) - \frac{f(t_{n-1}, y_{n-1})}{h}(t - t_n)\right] dt \end{aligned} \tag{51}$$

Simplyfying further, we get

$$\begin{aligned} y(t_{n+1}) &= y(t_n) + \frac{1-\alpha}{\mathbb{M}(\alpha)}[f(t_n, y(t_n)) - f(t_{n-1}, y(t_{n-1}))] \\ &+ \frac{\alpha}{\mathbb{M}(\alpha)} \left[\frac{f(t_n, y_n)}{h}\left[\frac{t_{n+1}^2}{2} - t_{n+1}t_{n-1} - \frac{t_n^2}{2} + t_n t_{n-1}\right] - \frac{f(t_{n-1}, y_{n-1})}{h}\left[\frac{t_{n+1}^2}{2} - t_{n+1}t_n + \frac{t_n^2}{2}\right]\right] \end{aligned} \tag{52}$$

Substituting $h = t_n - t_{n-1} = t_{n+1} - t_n$ in (52), we have

$$\begin{aligned} y(t_{n+1}) &= y(t_n) + \frac{1-\alpha}{\mathbb{M}(\alpha)}[f(t_n, y(t_n)) - f(t_{n-1}, y(t_{n-1}))] \\ &+ \frac{\alpha}{\mathbb{M}(\alpha)} \left[\frac{3h}{2} f(t_n, y_n) - \frac{h}{2} f(t_{n-1}, y_{n-1})\right] \end{aligned} \tag{53}$$

Thus, we arrive at an approximate solution by rearranging the eqn (53) and get

$$y(t_{n+1}) = y(t_n) + \left(\frac{1-\alpha}{M(\alpha)} + \frac{3\alpha h}{2M(\alpha)}\right) f(t_n, y(t_n)) - \left(\frac{1-\alpha}{M(\alpha)} + \frac{\alpha h}{2M(\alpha)}\right) f(t_{n-1}, y(t_{n-1})) \tag{54}$$

**Remark S5.** *Substituting $\alpha = 1$ in eqn (54) and simplifying, we obtain*

$$y(t_{n+1}) = y(t_n) + \frac{3}{2} h f(t_n, y_n) - \frac{1}{2} h f(t_{n-1}, y_{n-1}) \tag{55}$$

*which essentially represents the classical Adams-Bashforth scheme.*

**Remark S6.** *The approximate numerical solution obtained (eqn 43) for Caputo - Fabrizio FIVP via the Adams-Bashforth scheme is apparently different from eqn (3.26) of Atangana and Owolabi (2018).*

*S2.1.3. Atangana-Baleanu-Caputo fractional derivative*

Consider the general FDE with decaying memory in the form of the Mittag-Leffler function as

$$\begin{aligned} {}_0^{ABC}D_t^{\alpha} y(t) &= f(t, y(t)),\ \alpha \in (0,1], \\ y(0) &= y_0. \end{aligned} \tag{56}$$

By taking inverse Laplace transform and using the convolution theorem (Atangana, 2016) on eqn (56), we get

$$y(t) - y(0) = \frac{1-\alpha}{\mathbb{ABC}(\alpha)}[f(t, y(t))] + \frac{\alpha}{\mathbb{ABC}(\alpha)\Gamma(\alpha)} \int_0^t [(t-k)^{(\alpha-1)} f(k, y(k))]\, dk. \tag{57}$$

So, at $t = t_{n+1}$, where $n = 0, 1, 2, ...$, we obtain

$$y(t_{n+1}) - y(0) = \frac{1-\alpha}{\mathbb{ABC}(\alpha)}[f(t_n, y(t_n))] + \frac{\alpha}{\mathbb{ABC}(\alpha)\Gamma(\alpha)} \int_0^{t_{n+1}} [(t_{n+1}-t)^{(\alpha-1)} f(t, y(t))]\, dt \tag{58}$$

and at $t = t_n$

$$y(t_n) - y(0) = \frac{1-\alpha}{\mathbb{ABC}(\alpha)}[f(t_{n-1}, y(t_{n-1}))] + \frac{\alpha}{\mathbb{ABC}(\alpha)\Gamma(\alpha)} \int_0^{t_n} [(t_n-t)^{(\alpha-1)} f(t, y(t))]\, dt \tag{59}$$

Subtracting eqn. (59) from eqn. (58) yields

$$\begin{aligned} y(t_{n+1}) &= y(t_n) + \frac{1-\alpha}{\mathbb{ABC}(\alpha)}\left[f(t_n, y(t_n)) - f(t_{n-1}, y(t_{n-1}))\right] \\ &\quad + \frac{\alpha}{\mathbb{ABC}(\alpha)\Gamma(\alpha)} \int_0^{t_{n+1}} [(t_{n+1}-t)^{(\alpha-1)} f(t, y(t))]\, dt - \frac{\alpha}{\mathbb{ABC}(\alpha)\Gamma(\alpha)} \int_0^{t_n} [(t_n-t)^{(\alpha-1)} f(t, y(t))]\, dt \end{aligned} \tag{60}$$

For simplicity, we will use the form

$$y(t_{n+1}) = y(t_n) + \frac{1-\alpha}{\mathbb{ABC}(\alpha)}\left[f(t_n, y(t_n)) - f(t_{n-1}, y(t_{n-1}))\right] + AB_1 - AB_2 \tag{61}$$

where

$$AB_1 = \frac{\alpha}{\mathbb{ABC}(\alpha)\Gamma(\alpha)} \int_0^{t_{n+1}} [(t_{n+1}-t)^{(\alpha-1)} f(t, y(t))]\, dt \tag{62}$$

and

$$AB_2 = \frac{\alpha}{\mathbb{ABC}(\alpha)\Gamma(\alpha)} \int_0^{t_n} [(t_n-t)^{(\alpha-1)} f(t, y(t))]\, dt \tag{63}$$

Upon approximating the $f(t, y(t))$ using the Lagrange interpolation and proceeding in same manner as in Caputo fractional derivative discretization (34), we get where

$$\begin{aligned} AB_1 &= \frac{f(t_n, y_n)}{h\mathbb{ABC}(\alpha)\Gamma(\alpha)}\left[\frac{t_{n+1}^{\alpha+1}}{(\alpha+1)} - t_{n+1}^{\alpha} t_{n-1}\right] - \frac{f(t_{n-1}, y_{n-1})}{h\mathbb{ABC}(\alpha)\Gamma(\alpha)}\left[\frac{t_{n+1}^{\alpha+1}}{(\alpha+1)} - t_{n+1}^{\alpha} t_n\right] \\ AB_2 &= \frac{f(t_n, y_n)}{h\mathbb{ABC}(\alpha)\Gamma(\alpha)}\left[\frac{t_n^{\alpha+1}}{(\alpha+1)} - t_n^{\alpha} t_{n-1}\right] - \frac{f(t_{n-1}, y_{n-1})}{h\mathbb{ABC}(\alpha)\Gamma(\alpha)}\left[\frac{\alpha\, t_n^{\alpha+1}}{(\alpha+1)}\right] \end{aligned} \tag{64}$$

**Remark S7.** *The term $AB_2$ in equation (64) is different when compared to the initially solved analogous term $A_{\alpha,2}$ (eqn 3.37) in Atangana and Owolabi (2018) and in the revised version (Atangana and Owolabi, 2021).*

Thus, we arrive at approximate solution by by substituting $AB_1$ and $AB_2$ in (61) and get

$$\begin{aligned} y(t_{n+1}) &= y(t_n) + \frac{1-\alpha}{\mathbb{ABC}(\alpha)}\left[f(t_n, y(t_n)) - f(t_{n-1}, y(t_{n-1}))\right] \\ &+ \frac{f(t_n, y(t_n))}{\mathbb{ABC}(\alpha)\Gamma(\alpha)h}\left[\frac{t_{n+1}^{\alpha+1}}{(\alpha+1)} - t_{n+1}^{\alpha}t_{n-1} - \frac{t_n^{\alpha+1}}{(\alpha+1)} + t_n^{\alpha}t_{n-1}\right] \\ &- \frac{\alpha f(t_{n-1}, y(t_{n-1}))}{\mathbb{ABC}(\alpha)\Gamma(\alpha)h}\left[\frac{t_{n+1}^{\alpha+1}}{\alpha(\alpha+1)} - \frac{t_{n+1}^{\alpha}t_n}{\alpha} + \frac{t_n^{\alpha+1}}{\alpha+1}\right], \end{aligned} \tag{65}$$

**Remark S8.** *when $\alpha = 1$, eqn (65) will reduce to same form as (44), which straightforwardly reduces to classical Adams-Bashforth scheme.*

**Remark S9.** *The numerical discretization of Atangana-Baleanu-Caputo FIVP using Adams-Bashforth scheme and its application to chaotic problems is also discussed in Owolabi and Atangana (2019), but the article was further retracted due to unreliability in extending local approximation method to non-local problems (Owolabi and Atangana, 2021).*

#### *S2.2. Riemann-Liouville type FDEs*

FDEs with Riemann-Liouville fractional operator (R-L (S1.2), CFR (S1.7)) have been approximated approximated using the efficient finite difference numerical technique (Baleanu et al., 2012). The initial exploration of approximate solutions for FDEs with the Riemann-Liouville fractional operator was conducted by Atangana and Gómez-Aguilar (2018) and subsequent work by Owolabi (2018) discussed the application of approximation to chaotic systems. However, we identified minor calculation errors in the numerical derivation (Atangana and Gómez-Aguilar, 2018), and a modified final form is adopted for likelihood computation. Hence, we opted to re-derive the numerical scheme, in short, to maintain a coherent flow throughout the article. The stability and convergence results of the approximate solutions are discussed in Atangana and Gómez-Aguilar (2018) and Owolabi (2018).

##### *S2.2.1. Riemann-Liouville FDE with power-law decay*

Consider a general logistic FDE of the form

$$\begin{aligned} {}_0^{RL}D_t^{\alpha} f(t) &= F(t, f(t)), \\ f(0) &= f_0. \end{aligned} \tag{66}$$

Using eqn. (4) on Left Hand Side (LHS) of (66), we obtain

$$\begin{aligned} {}_0^{RL}D_t^{\alpha}\{f(t)\} &= \frac{1}{\Gamma(1-\alpha)}\frac{d}{dt}\int_0^t f(k)(t-k)^{-\alpha}\,dk, \quad \alpha \in (0, 1], \\ {}_0^{RL}D_t^{\alpha}\{f(t)\} &= \frac{d}{dt}u(t) \end{aligned} \tag{67}$$

where $f(0) = f_0$ and $u(t) = \frac{1}{\Gamma(1-\alpha)}\int_0^t f(k)(t-k)^{-\alpha}$.

Applying the forward approximation method on the Right Hand Side (RHS) of eqn. (67), we get

$$\frac{d}{dt}u(t) = \frac{u(t_{j+1}) - u(t_j)}{h} \tag{68}$$

where,

$$u(t_{j+1}) = \frac{1}{\Gamma(1-\alpha)}\int_0^{t_{j+1}} f(k)(t_{j+1}-k)^{-\alpha}\,dk, \tag{69}$$

and

$$u(t_j) = \frac{1}{\Gamma(1-\alpha)}\int_0^{t_j} f(k)(t_j-k)^{-\alpha}\,dk. \tag{70}$$

Upon using trapezoidal approximation rule on $u(t_{j+1})$ (69) and ignoring the approximation error, we obtain

$$\begin{aligned} u(t_{j+1}) &= \frac{1}{\Gamma(1-\alpha)}\sum_{s=0}^{j}\int_{t_s}^{t_{s+1}} \frac{f(t_{s+1})+f(t_s)}{2}(t_{j+1}-k)^{-\alpha}\,dk \\ &= \frac{1}{\Gamma(2-\alpha)}\sum_{s=0}^{j}\frac{f(t_{s+1})+f(t_s)}{2}\left[(t_{j+1}-t_s)^{1-\alpha}-(t_{j+1}-t_{s+1})^{1-\alpha}\right] \end{aligned} \tag{71}$$

**Remark S10.** *The term $u(t_{j+1})$ in equation (71) differs when compared to the initially solved analogous term $u(t_{j+1})$ (eqn 16) in Owolabi (2018) and $F(t_{j+1})$(eqn 7) in Atangana and Gómez-Aguilar (2018).*

Similarly,

$$u(t_j) = \frac{1}{\Gamma(2-\alpha)}\sum_{s=0}^{j-1}\frac{f(t_{s+1})+f(t_s)}{2}\left[(t_j-t_s)^{1-\alpha}-(t_j-t_{s+1})^{1-\alpha}\right] \tag{72}$$

Substituting $u(t_{j+1})$ (71) and $u(t_j)$ (72) in eqn. (68), we have

$$\begin{aligned} \frac{d}{dt}u(t) &= \frac{1}{h\Gamma(2-\alpha)}\left[\sum_{s=0}^{j}\frac{f(t_{s+1})+f(t_s)}{2}\left[(t_{j+1}-t_s)^{1-\alpha}-(t_{j+1}-t_{s+1})^{1-\alpha}\right]\right] \\ &\quad -\frac{1}{h\Gamma(2-\alpha)}\left[\sum_{s=0}^{j-1}\frac{f(t_{s+1})+f(t_s)}{2}\left[(t_j-t_s)^{1-\alpha}-(t_j-t_{s+1})^{1-\alpha}\right]\right] \end{aligned} \tag{73}$$

Thus, upon further simplifying and using (66), the approximate solution is

$$\begin{aligned} f(t_{j+1}) &= -f(t_j) - h^{\alpha-1}\sum_{s=0}^{i-1}(f(t_{s+1})+f(t_s))\Big[(t_{j+1}-t_s)^{(1-\alpha)} \\ &\quad -(t_{j+1}-t_{s+1})^{(1-\alpha)}-(t_j-t_s)^{(1-\alpha)}+(t_j-t_{s+1})^{(1-\alpha)}\Big] \\ &\quad +2h^{\alpha}\Gamma(2-\alpha)F(t_j,f(t_j)) \end{aligned} \tag{74}$$

*S2.2.2. Riemann-Liouville FDE with exponential decay law*

Consider a general logistic FDE with exponential decay of the form

$$\begin{aligned} {}_0^{CFR}D_t^{\alpha}f(t) &= F(t,f(t)), \quad \alpha\in(0,1], \\ f(0) &= f_0. \end{aligned} \tag{75}$$

Using eqn. (22) on Left Hand Side (LHS) of (75), we obtain

$$
\begin{aligned}
{}_{0}^{RL}D_t^{\alpha}\{f(t)\} &= \frac{\mathbb{M}(\alpha)}{(1-\alpha)}\frac{d}{dt}\int_0^t f(k)\exp\left(\frac{-\alpha(t-k)}{(1-\alpha)}\right)dk, \\
{}_{0}^{RL}D_t^{\alpha}\{f(t)\} &= \frac{d}{dt}u(t)
\end{aligned} \tag{76}
$$

where $f(0) = f_0$ and $u(t) = \frac{\mathbb{M}(\alpha)}{(1-\alpha)}\int_0^t f(k)\exp\left(\frac{-\alpha(t-k)}{(1-\alpha)}\right)dk$.

Proceeding in the same manner as in from eqn. (67) to (70), we get

$$
\begin{aligned}
u(t_{j+1}) &= \frac{\mathbb{M}(\alpha)}{(1-\alpha)}\sum_{s=0}^{j}\int_{t_s}^{t_{s+1}}\frac{f(t_{s+1})+f(t_s)}{2}\exp\left(\frac{-\alpha(t_{j+1}-k)}{(1-\alpha)}\right)dk \\
&= \frac{\mathbb{M}(\alpha)}{\alpha}\sum_{s=0}^{j}\frac{f(t_{s+1})+f(t_s)}{2}\left[\exp\left(\frac{-\alpha(t_{j+1}-t_{s+1})}{(1-\alpha)}\right)-\exp\left(\frac{-\alpha(t_{j+1}-t_s)}{(1-\alpha)}\right)\right]
\end{aligned} \tag{77}
$$

Similarly,

$$
u(t_j) = \frac{\mathbb{M}(\alpha)}{\alpha}\sum_{s=0}^{j-1}\frac{f(t_{s+1})+f(t_s)}{2}\left[\exp\left(\frac{-\alpha(t_j-t_{s+1})}{(1-\alpha)}\right)-\exp\left(\frac{-\alpha(t_j-t_s)}{(1-\alpha)}\right)\right] \tag{78}
$$

Substituting $u(t_{j+1})$ (77) and $u(t_j)$ (78) in eqn. (76), we have

$$
\begin{aligned}
\frac{d}{dt}u(t) &= \frac{\mathbb{M}(\alpha)}{h\alpha}\sum_{s=0}^{j}\frac{f(t_{s+1})+f(t_s)}{2}\left[\exp\left(\frac{-\alpha(t_{j+1}-t_{s+1})}{(1-\alpha)}\right)-\exp\left(\frac{-\alpha(t_{j+1}-t_s)}{(1-\alpha)}\right)\right] \\
&\quad -\frac{\mathbb{M}(\alpha)}{h\alpha}\sum_{s=0}^{j-1}\frac{f(t_{s+1})+f(t_s)}{2}\left[\exp\left(\frac{-\alpha(t_j-t_{s+1})}{(1-\alpha)}\right)-\exp\left(\frac{-\alpha(t_j-t_s)}{(1-\alpha)}\right)\right]
\end{aligned} \tag{79}
$$

**Remark S11.** *Eqn.(79) differs when compared to the the originally solved counterpart in Atangana and Gómez-Aguilar (2018), despite our replacement of $\sum_{s=0}^{j-1}$ with $\sum_{s=1}^{j}$ and corresponding adjustment of second term of equation (79).*

Thus, upon further simplifying and using (75), the approximate solution is

$$
\begin{aligned}
f(t_{j+1}) &= -f(t_j) + \frac{1}{\left(1-\exp\left(\frac{-\alpha h}{1-\alpha}\right)\right)}\sum_{s=0}^{j-1}(f(t_{s+1})+f(t_s)) \times \\
&\quad \left[\exp\left(\frac{-\alpha}{1-\alpha}(t_j-t_{s+1})\right)-\exp\left(\frac{-\alpha}{1-\alpha}(t_j-t_s)\right)-\right. \\
&\quad \left.\exp\left(\frac{-\alpha}{1-\alpha}(t_{j+1}-t_{s+1})\right)+\exp\left(\frac{-\alpha}{1-\alpha}(t_{j+1}-t_s)\right)\right] \\
&\quad +\frac{2\alpha h}{M(\alpha)\left(1-\exp\left(\frac{-\alpha h}{1-\alpha}\right)\right)}F(t_j, f(t_j))
\end{aligned} \tag{80}
$$

#### S2.3. Grunwald and Letnikov FDEs

FDEs defined Grunwald - Letnikov sense have a discrete structure hence the numerical discretization was done using simple taylor's approximation as in Ariza-Hernandez et al. (2020b). The approximate numerical solution of a general FIVP where fractional operator is defined in G-L sense is

$$f(t_{j+1}) = \sum_{s=1}^{j+1}(-1)^{s+1}\binom{\alpha}{s}f(t_{j+1-s}) + h^{\alpha}F(t_j, f(t_j)) \tag{81}$$

#### S2.4. Numerically approximated fractional logistic model

The final numerically discretized solution of fractional logistic model where $\mu^{(m)}(t_i)$ represents the solution for $i \in 1, 2, \cdots, n-1$ discrete time points and $m \in 1, 2, \cdots, 6$ corresponds to each definition of FDs is

$$\begin{aligned}\mu^{(1)}(t_i) &= N(t_i) + \frac{rN(t_i)\left(1 - \frac{N(t_i)}{K}\right)}{h\Gamma(\alpha)}\left[\frac{t_{i+1}^{\alpha+1}}{\alpha(\alpha+1)} - \frac{t_{i+1}^{\alpha}t_{i-1}}{\alpha} - \frac{t_i^{\alpha+1}}{\alpha(\alpha+1)} + \frac{t_i^{\alpha}t_{i-1}}{\alpha}\right] \\ &\quad - \frac{rN(t_{i-1})\left(1 - \frac{N(t_{i-1})}{K}\right)}{h\Gamma(\alpha)}\left[\frac{t_{i+1}^{\alpha+1}}{\alpha(\alpha+1)} - \frac{t_{i+1}^{\alpha}t_i}{\alpha} + \frac{t_i^{\alpha+1}}{\alpha+1}\right],\end{aligned} \tag{82}$$

$$\begin{aligned}\mu^{(2)}(t_i) &= N(t_i) + \left(\frac{1-\alpha}{M(\alpha)} + \frac{3\alpha h}{2M(\alpha)}\right)rN(t_i)\left(1 - \frac{N(t_i)}{K}\right) \\ &\quad - \left(\frac{1-\alpha}{M(\alpha)} + \frac{\alpha h}{2M(\alpha)}\right)rN(t_{i-1})\left(1 - \frac{N(t_{i-1})}{K}\right),\end{aligned} \tag{83}$$

$$\begin{aligned}\mu^{(3)}(t_i) &= N(t_i) + \frac{1-\alpha}{ABC(\alpha)}\left[rN(t_i)\left(1 - \frac{N(t_i)}{K}\right) - rN(t_{i-1})\left(1 - \frac{N(t_{i-1})}{K}\right)\right] \\ &\quad + \frac{\alpha rN(t_i)\left(1 - \frac{N(t_i)}{K}\right)}{ABC(\alpha)\Gamma(\alpha)h}\left[\frac{t_{i+1}^{\alpha+1}}{\alpha(\alpha+1)} - \frac{t_{i+1}^{\alpha}t_{i-1}}{\alpha} - \frac{t_i^{\alpha+1}}{\alpha(\alpha+1)} + \frac{t_i^{\alpha}t_{i-1}}{\alpha}\right] \\ &\quad - \frac{\alpha rN(t_{i-1})\left(1 - \frac{N(t_{i-1})}{K}\right)}{ABC(\alpha)\Gamma(\alpha)h}\left[\frac{t_{i+1}^{\alpha+1}}{\alpha(\alpha+1)} - \frac{t_{i+1}^{\alpha}t_i}{\alpha} + \frac{t_i^{\alpha+1}}{\alpha+1}\right],\end{aligned} \tag{84}$$

$$\mu^{(4)}(t_i) = -N(t_i) - h^{\alpha-1}\sum_{s=0}^{i-1}\left(N(t_{s+1}) + N(t_s)\right)\Big[(t_{i+1}-t_s)^{(1-\alpha)} - (t_{i+1}-t_{s+1})^{(1-\alpha)} - (t_i - t_s)^{(1-\alpha)} + (t_i - t_{s+1})^{(1-\alpha)}\Big] + 2h^{\alpha}\Gamma(2-\alpha) r N(t_i)\left(1 - \frac{N(t_i)}{K}\right), \tag{85}$$

$$\mu^{(5)}(t_i) = -N(t_i) + \frac{1}{\left(1-\exp\left(\frac{-\alpha h}{1-\alpha}\right)\right)}\sum_{k=0}^{i-1}(N(t_{k+1}) + N(t_k)) \times \Big[\exp\left(\frac{-\alpha}{1-\alpha}(t_i - t_{k+1})\right) - \exp\left(\frac{-\alpha}{1-\alpha}(t_i - t_k)\right) - \exp\left(\frac{-\alpha}{1-\alpha}(t_{i+1} - t_{k+1})\right) + \exp\left(\frac{-\alpha}{1-\alpha}(t_{i+1} - t_k)\right)\Big] + \frac{2\alpha h}{M(\alpha)\left(1-\exp\left(\frac{-\alpha h}{1-\alpha}\right)\right)} r N(t_i)\left(1 - \frac{N(t_i)}{K}\right), \tag{86}$$

$$\mu^{(6)}(t_i) = \sum_{s=1}^{i+1}(-1)^{s+1}\binom{\alpha}{s} N(t_{i+1-s}) + h^{\alpha} r N(t_i)\left(1 - \frac{N(t_i)}{K}\right). \tag{87}$$

### S3. Parameter Distributions of Alternative Models: Real Data Analysis

The Density plots for the model parameters $\alpha$, $r, K$ and $\tau = 1/\sigma^2$ of the most suitable model $M_6$ (GF FD) are reported in the main text. Rest of the models ($M_1$ to $M_5$) parameters density plots are depicted in the following figures (S1-S4) for the *Ursus Americanus* (GPDD ID: 116), *Castor canadensis* (GPDD ID: 200), *Phalacrocorax carbo* (GPDD ID: 9330) and Covid-19 total cases data for Germany, respectively.

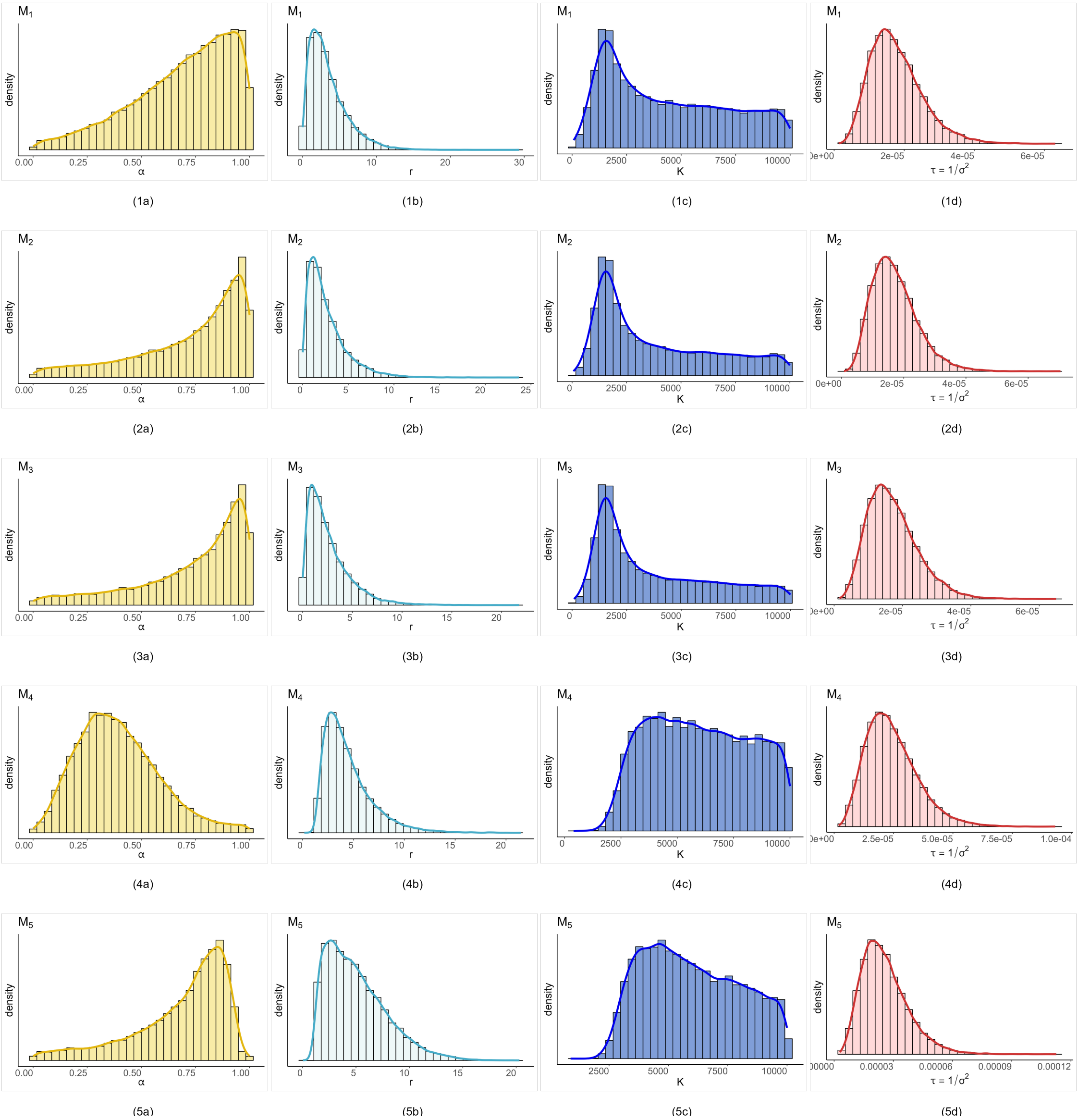

Figure S1: Density plots for model parameters $\alpha,\ r, K$ and $\tau = 1/\sigma^2$ of models $M_1$ to $M_5$ (Caputo FD to CF-RL FD) corresponds to the data from GPDD of species *Ursus Americanus* (GPDD ID: 116). The posterior distribution is obtained using `JAGS`, and the overlaid curve of posterior density is approximated using the Gaussian kernel density estimator.

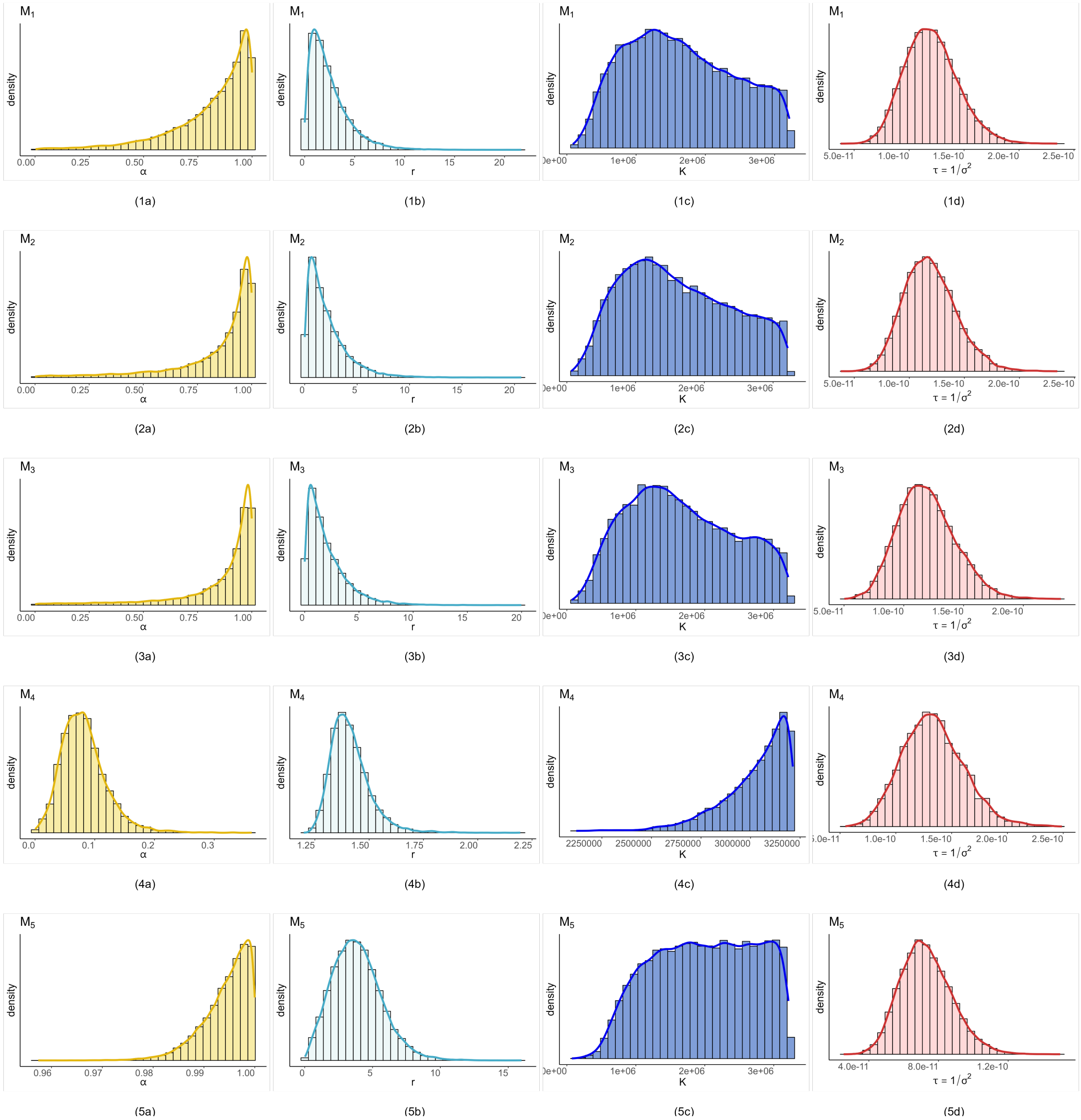

Figure S2: Density plots for model parameters $\alpha,\ r, K$ and $\tau = 1/\sigma^2$ of model $M_1$ to $M_5$ (Caputo FD to CF-RL FD) corresponds to the data from GPDD of species *Castor canadensis* (GPDD ID: 200). The posterior distribution is obtained using `JAGS`, and the overlaid curve of posterior density is approximated using the Gaussian kernel density estimator.

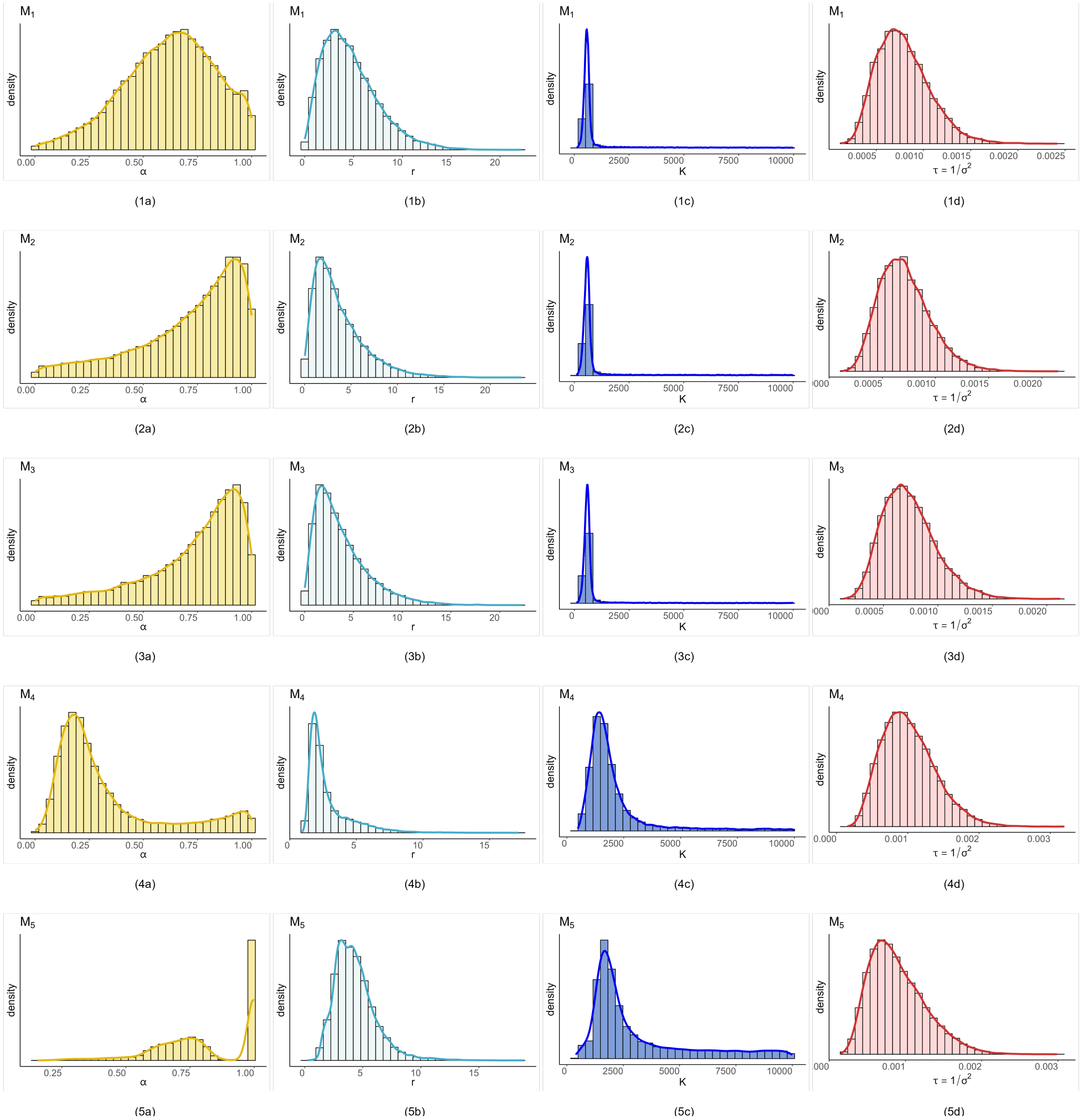

Figure S3: Density plots for model parameters $\alpha,\ r, K$ and $\tau = 1/\sigma^2$ of model $M_1$ to $M_5$ (Caputo FD to CF-RL FD) corresponds to the data from GPDD of species *Phalacrocorax carbo* (GPDD ID: 9330). The posterior distribution is obtained using `JAGS`, and the overlaid curve of posterior density is approximated using the Gaussian kernel density estimator.

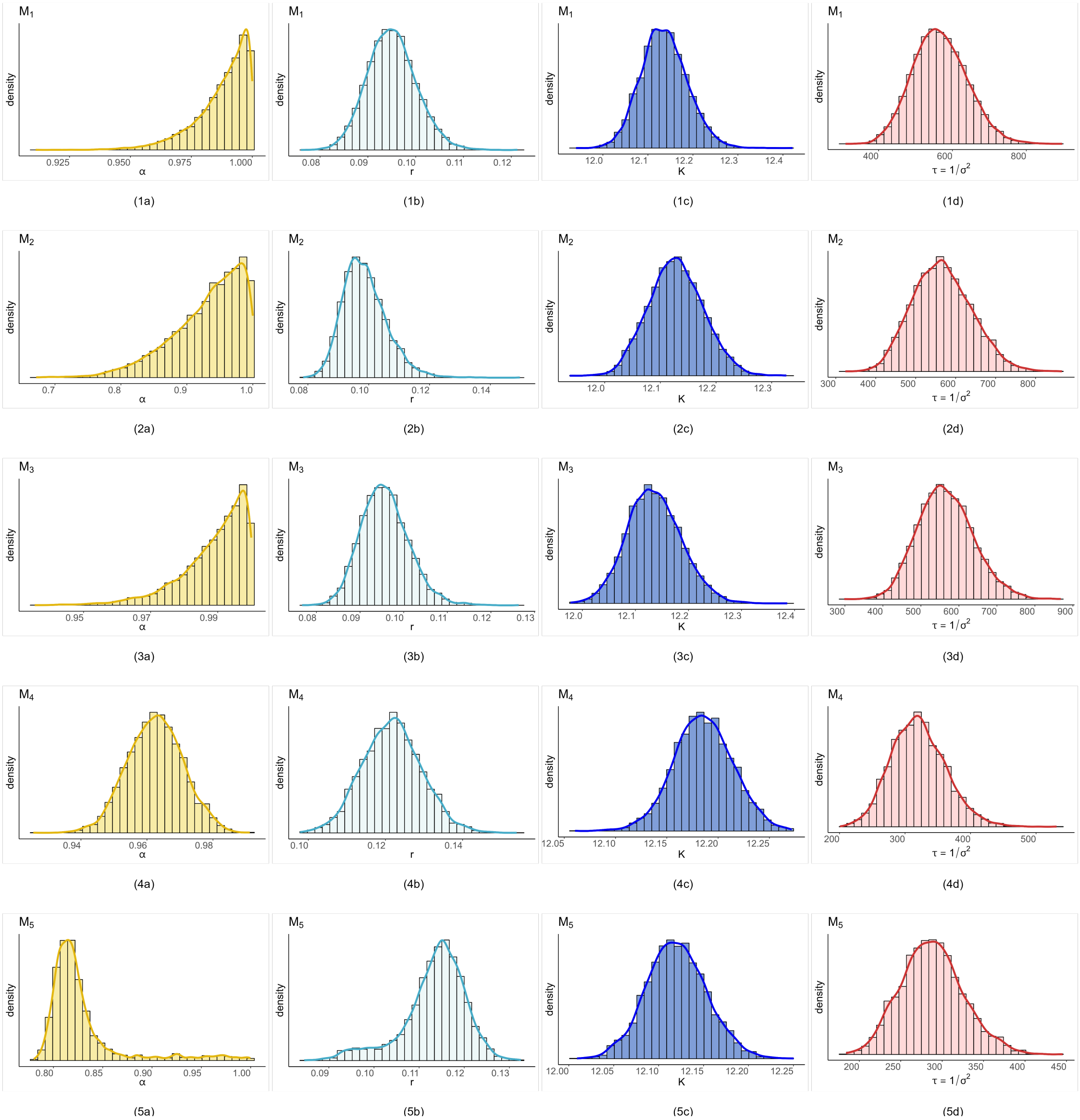

Figure S4: Density plots for model parameters $\alpha,\ r, K$ and $\tau = 1/\sigma^2$ of model $M_1$ to $M_5$ (Caputo FD to CF-RL FD) corresponds to Covid-19 confirmed total cases data (natural logarithmic scale) for Germany. The posterior distribution is obtained using `JAGS`, and the overlaid curve of posterior density is approximated using the Gaussian kernel density estimator.

### S4. Fractional Differential definition selection (R codes)

1. All the analyses are done using the `R` software.
2. The codes for reproducing analyses, figures, and applying used methodology are freely available at `https://github.com/dipalimestry96/BaFOMS-Bayesian-Fractional-Order-Model-Selection`.
3. Codes are in the folder named "Data Analysis Mestry et al 2024". To access them, refer to the folder hierarchy given in Figure S5.

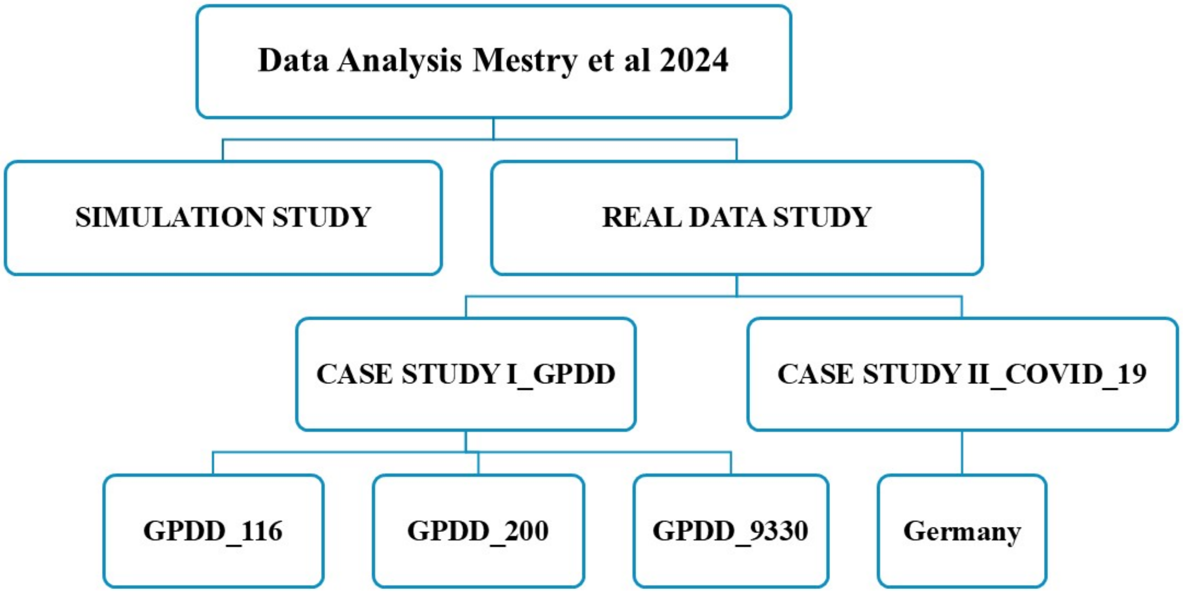

Figure S5: All the code files used in this study are organized within the main folder titled “Data Analysis Mestry et al. 2024”. This folder is divided into two sub-folders: “SIMULATION STUDY” for simulated data analysis and “REAL DATA STUDY” for real data analysis. The analysis includes two primary datasets, the Global Population Dynamics Database (GPDD) and COVID-19, with corresponding code files located in the “CASE STUDY I_GPDD” and “CASE STUDY II_COVID_19” folders, respectively. Within the GPDD case study, three species datasets—*Ursus americanus* (GPDD ID:116), *Castor canadensis* (GPDD ID:200), and *Phalacrocorax carbo* (GPDD ID:9330) are analyzed, with relevant code files stored in the sub-folders “GPDD_116”, “GPDD_200”, and “GPDD_9330”. For the COVID-19 case study, data for Germany is analyzed, with the corresponding code files located in the “Germany” sub-folder within “CASE STUDY II_COVID-19”.

Zeng, Fanhai, C.L., 2015. Numerical Methods for Fractional Calculus. Chapman and Hall/CRC, New York.
